# Hemodynamic and electrophysiological progression of the Rose Bengal photothrombotic stroke model in mice: vasoactive properties of Rose Bengal, tissue heating, wavelength optimization, and sex differences in lesion volume

**DOI:** 10.1101/2025.09.22.677889

**Authors:** Patrick Chary, Sarah Rehmani, Simone Davidson, Xiaonan Li, Simon X. Chen, Gergely Silasi

## Abstract

The photothrombotic stroke model is gaining popularity due to its relative simplicity, minimal invasiveness, and clinical relevance. Photothrombosis involves the delivery of an intravascular photosensitizer (Rose Bengal) followed by its photoactivation, resulting in vessel occlusion and ischemia. Using a combination of complementary optical and non-optical techniques, we characterized the physiological changes in mice undergoing photothrombosis. We report that Rose Bengal acts as a rapid vasoconstrictor, inducing hypoemia both in the brain and periphery even in the absence of its photoactivation. Conversely, we find that light, when used at photothrombosis-appropriate intensities and durations, induces large amounts of tissue heating and hyperemia even in the distal non-illuminated hemisphere. Furthermore, we show that use of the optimal photothrombotic wavelength based on the Rose Bengal absorption spectrum (yellow-561nm) produces a more consistent and pronounced drop in blood flow, and a shorter latency to the initial spreading depolarization (SD), ultimately resulting in a larger stroke. Similarly, when yellow light is used to induce a stroke in ChR2-expressing mice, the electrophysiological and hemodynamic confounds from green light cross activation of ChR2 are eliminated. Finally, we observe across cohorts that male mice have larger strokes than females. Altogether, we extensively describe important caveats and confounds concerning photothrombosis and provide a detailed characterization of its early ischemic events.

**Significance statement:** Photothrombosis is a powerful model of ischemic stroke which uses light to photoactivate an intravascular dye (Rose Bengal). However, little is known about the independent effects of both the Rose Bengal and the light used to activate it. We show that both manipulations introduce separate confounds relevant to stroke outcomes, something which should be considered and accounted for when using this technique. In addition, we demonstrate that by using the optimal Rose Bengal excitation wavelength, the blood flow drop is more pronounced and consistent, resulting in larger strokes and perhaps better modelling human injury. Furthermore, we show that precautions can be taken to avoid spectral overlap when integrating photothrombosis in optogenetic experiments. Finally, we explore sex differences in lesion volume.

## Introduction

Animal models of ischemic stroke are diverse and there is no gold standard most relevant to human disease (Mergenthaler and Meisel, 2012; Corbett et al., 2017). Photothrombosis has emerged as a popular choice because it is relatively simple to perform, minimally invasive, possible in behaving animals, highly controllable with regards to ischemia location, and relatively representative of ischemic stroke in humans (Ginsberg and Busto, 1989; Carmichael, 2005; Schmidt et al., 2012; Labat-gest and Tomasi, 2013; Fluri et al., 2015; Uzdensky, 2018). Photothrombosis happens in two parts, requiring the delivery of an intravascular photosensitizer followed by its photoactivation. The photochemistry produces reactive oxygen species, mostly singlet oxygen via type II photosensitized oxidation reactions (Baptista et al., 2017), which damage the inside of blood vessels and thrombose them via recruitment of members from the clotting cascade (Rosenblum and El-Sabban, 1977; Watson et al., 1985; Matsuno et al., 1993). Historically, several photosensitizers have been used for photothrombotic purposes, including Erythrosin B (Wester et al., 1995; Hilger et al., 2004), Indocyanine Green (Costa et al., 2002; Navajas et al., 2003; Farah et al., 2004; Arevalo et al., 2005), Photofrin (Futrell, 1991; Ishikawa et al., 2002), Phthalocyanine (Pallikaris et al., 1993), Verteporfin (Yoon et al., 2007), and Fluorescein (Rosenblum and El-Sabban, 1977; Herrmann, 1983). However, the photosensitizer which has come to dominate photothrombosis protocols is Rose Bengal (RB), likely owing to a combination of its high water solubility, high singlet oxygen quantum yield, low rate of photodegradation, and ease of sourcing (Neckers, 1989; Alarcón et al., 2009). Perhaps because of these advantages, the model was rapidly and extensively adapted in rodent studies, however a detailed description of the physiological effects of the individual components of the model (RB and light) has not been carried out.

Measures of stroke outcome such as lesion volume, behavioral deficits and functional connectivity are known to be affected by the physiological state of an animal preceding, during, and immediately after injury. For instance, both ischemic pre-conditioning (Weir et al., 2021) and mild post-stroke hypothermia (Busto et al., 1989; Zhang et al., 2022; Lyu et al., 2025) are neuroprotective, while hyperthermia (Busto et al., 1987; Kim et al., 1996; Reglodi et al., 2000; Sinigaglia-Coimbra et al., 2002) worsens outcomes. Careful measurements of blood flow have shown that the magnitude and inter-animal consistency of induced ischemia during stroke directly predicts the severity of injury (Wang et al., 2022; Marks et al., 2025). Similarly, the quantity and origin of spreading depolarizations (SDs) after stroke are important predictors of stroke severity as their occurrence can impact histological measures as well as behavioural outcomes (Risher et al., 2010; von Bornstädt et al., 2015; Schoknecht et al., 2021; Boyce et al., 2025). Altogether, care must be taken to understand how physiological parameters such as blood flow, brain temperature and large-scale electrical activity are altered during stroke, but also by the procedures used to induce the ischemia.

We report that RB reduces cerebral as well as peripheral blood flow even in the absence of its photoactivation, likely through its action as a rapid vasoconstrictor. Conversely, we show that on its own, the light used for RB photoactivation causes both significant increases in cortical blood flow and tissue heating, even in the non-illuminated hemisphere. Guided by the optimal absorption spectrum of RB under physiological conditions, we show that activation of RB by yellow light induces more severe ischemia, reduces the onset of SDs, and produces larger lesions relative to green light. We show that green light can inadvertently induce aberrant cortical activity in ChR2-expressing mice as a result of spectral overlap, which is critical for many experiments that employ ChR2 for therapeutic stimulation or for probing specific circuits after PT stroke. Finally, we describe the large-scale changes in electrical activity and blood flow during progression of the photothrombotic stroke and show that male mice have larger lesions than female mice.

## Materials and methods

### Animals

Mice were bred and housed in a 12h light/12h dark cycle. The mice had *ad libitum* access to chow and water. All experimental procedures were conducted in accordance with the guidelines of the Canadian Council on Animal Care. Experimental protocols were approved by the University of Ottawa Animal Care Committee.

The opsin expressing mouse strain used was the Thy1-ChR2-YFP (B6. Cg-Tg(Thy1-COP4/EYFP)18Gfng/J; 007612, The Jackson Laboratory) (Arenkiel et al., 2007; Wang et al., 2007; Fenno et al., 2015). This mouse strain expresses ChR2 in layer 5 cortical neurons. The mice are viable, fertile, normal in size and do not display any gross physical or behavioral abnormalities. The non-opsin expressing mouse strain used throughout this study was the Tmem119-2A-EGFP (C57BL/6-Tmem119em2Gfng/J; 031823, The Jackson Laboratory) (Kaiser and Feng, 2019), which expresses GFP in the microglia, allowing for their labelling (a feature not used in this study). This strain has no reported gross phenotypic or behavioral abnormalities.

### Surgical implantation of the transcranial window

A chronic transcranial window was implanted a minimum of 7 days before experimental recordings using previously described protocols with slight modification (Silasi et al., 2013, 2016). Namely, the shaft of the stainless-steel acupuncture needle (DongBang acupuncture – 0.25mm x 15mm) used for recording EEG was first sanded rough to increase stability when embedded within the Metabond dental cement. In addition, to ensure the electrodes were accurately placed in the epidural space, the impedance to a subcutaneous reference electrode in the neck was measured during implantation using a handheld LCR meter (Model: ET430; Amazon). The electrode was manually advanced into the bone until a resistance of less than 15kohms was measured. The clear version of C&B Metabond dental cement was prepared by mixing 1 scoop of C&B Metabond powder (Product: S399), 6 drops of C&B Metabond Quick Base (Product: S398) and one drop of C&B Universal catalyst (Product: S371) in a ceramic or glass dish (do not use plastic). Once the mixture reached a consistency that made it stick to the end of a wooden stir stick, a head-fixing screw was glued to the cerebellar plate by applying a small amount of dental cement to the butt end of a 4/40 stainless steel setscrew and holding it pressed against the skull until the cement partially dried. The screw was angled posteriorly (∼120° relative to skull) and was centered directly posterior of lambda. With the setscrew in place, a layer of dental adhesive was applied directly on the intact skull. A #1 cover glass (8mm ⌀ - Marienfeld) was gently placed on top of the mixture before it solidified (within 1 min), taking care to avoid bubble formation. If necessary, extra dental cement was applied around the edge of the cover slip to ensure that all exposed bone and the EEG electrodes were covered and that the incision site was sealed at the edges. The skin naturally tightened itself around the craniotomy and sutures were not necessary. Once the dental cement around the coverslip solidified completely (∼20 min), the animal was allowed to recover in the home cage.

### Laser doppler flowmetry and imaging (LDF and LDI)

LDI (Moor Instruments; moorLDI2-IR-HR) was used to generate both a baseline and post-stroke blood flow map by setting a scanning region over the entire chronic window (both hemispheres). LDF was used to continuously record blood flow from the site of photothrombosis before, during and after stroke induction. All laser doppler measurements were performed with a 2.5mW, 830nm laser, with a 40Hz sampling frequency. Control experiments were performed to ensure there was no crosstalk between the stimulation light (561nm) and the optical blood flow measurements (Fig.S1).

### Laser speckle contrast imaging (LSCI)

LSCI was performed by illuminating the chronic window with an 8mW, 830nm laser (model: M-16A830-250-X, Amazon) and recording the speckle pattern with a sCMOS camera (CS2100M-USB – ThorLabs) at 10Hz, 10ms exposure. Images were processed using a custom MATLAB (MathWorks) script to generate speckle pattern.

### Functional ultrasound

Transcranial functional ultrasound (fUS) imaging, a complementary hemodynamic monitoring tool to laser doppler and laser speckle, was also used to quantify RB induced changes in blood flow (Macé et al., 2011; Errico et al., 2015; Brunner et al., 2021). Mice were anesthetized with isoflurane, and the top of the head was shaved and treated with depilatory cream (Nair) to remove all hair. The mouse was placed in a stereotaxic frame and positioned under the fUS probe (IcoPrime; Iconeus). Body temperature was maintained at 37±1.5°C via a feedback-regulated heating pad and a rectal temperature probe. Ultrasound gel was applied generously and the fUS probe was carefully lowered over the scalp until a satisfactory image was produced. Images were generated at 2.5Hz before, during, and after both PBS and RB injections. PBS or RB were injected I.P (100mg/kg) into the prone mouse by lifting a hindleg.

### Craniotomy window and 2-photon vascular imaging

We used previously-developed methods for surgical implantation of the craniotomy window (Yin et al., 2021; Yang et al., 2022). Mice were subcutaneously injected with buprenorphine (0.05 mg/kg), Baytril (5 mg/kg), and Dexamethasone (2 mg/kg) as analgesia, anti-infection, and anti-inflammation, respectively and anesthetized with 1-2% of isoflurane. An incision was performed to remove a circular piece of the scalp; a custom head-plate was implanted on the skull with instant glue (Krazy Glue) and dental cement (Lang Dental). A craniotomy of approximately 2 mm in diameter was performed over the right motor cortex. A glass window was implanted over M1, and the implants were stabilized with Jet Denture Repair Powder Cement (Henry Schein). Mice recovered in the home cage for 3 weeks before any imaging.

On the day of imaging, FITC-Dextran 70 kDa (Sigma Aldrich – 46945-500MG-F) dissolved in PBS (50mg/mL) was injected (0.1mL) retro-orbitally (2% isoflurane) to label the cortical vasculature, before the anesthetized mouse was head-fixed underneath a commercial two-photon laser scanning microscope (Bergamo Scope, Thorlabs). Images were acquired at 30Hz through a 16x objective (NIKON) with excitation at 950 nm (InSight X3, Spectra-Physics). Body temperature was maintained at 37±1.5°C via a heating pad, and a winged infusion set (Terumo – SV-25BLK) filled with either PBS or RB (100mg/kg) was pre inserted into the I.P space. For each mouse, an imaging field containing many well resolved cortical vessels was selected. A pre injection Z-stack (though the entire column of visible vessels) was imaged and later made into a maximum intensity Z-projection. The plane with the most in focus vessels within this Z-stack was selected for time course imaging. A pre injection baseline of 26 seconds was recorded, then either PBS or RB was infused over 10 seconds, and the recording continued for an additional 100 seconds. Finally, a post injection Z-stack was acquired and made into a maximum intensity Z-projection. The mice were subsequently returned to the home cage and allowed to recover. Fluorescence intensity profiles for each in focus vessel were used to calculate the full width at half maximum (FWHM) using the VasoMetrics macro for ImageJ/Fiji (McDowell et al., 2021).

### Paw imaging

The ventral side of the mouse hind paw is relatively hairless and contains many surface blood vessels, making it amenable to non-invasive optical imaging of blood flow (Beringhs et al., 2020; Li et al., 2021; McDonald et al., 2021; Tang and Kim, 2021; Kam et al., 2023). Mice were anesthetized with isoflurane (5% induction and 2% maintenance in air), the eyes were treated with lubricant (Optimax) and the left hind limb was outstretched and taped down to a heating pad to immobilize during recording. A rectal temperature probe was inserted, and core temperature was maintained at 37±1.5°C via a feedback-regulated heating pad. LDI images of the ventral paw were taken before and after injections of either PBS or RB, and the LDF laser was positioned on the paw for the duration of the injections. PBS or RB (100mg/kg) was injected I.P into the prone mouse by lifting a hindleg. In our experience, blood flow in the paw is highly sensitive to even small fluctuations in core temperature; therefore, great care was taken to allow mice to reach a stable core temperature before initiating experiments.

### Thermal imaging

Measurements of brain temperature were made by thermal imaging through the transcranial window. Mice were anesthetized with isoflurane (5% induction and 2% maintenance in air) and head-fixed underneath a custom modular optical imaging system (LabeoTech – OiS200) via the cemented set screw and a locking ball and socket mount (Thorlabs, Product: TRB1). A rectal temperature probe was inserted, and core temperature was maintained at 37±1.5°C via a feedback-regulated heating pad. The eyes were treated with eye lubricant (Optixcare) to keep the corneas moist. The yellow light used for stimulation (561nm @ 7mW/mm^2^ @ 2.2mm ⌀ (DPSS – MGL-U-561 – Changchun New Industries Optoelectronics Technology Co)) was targeted 1.5mm lateral to bregma through the transcranial window via a system of galvo mirrors. A thermal camera (Waveshare electronics – MLX90640-D55) running on a Raspberry Pi 4 was positioned above the cranial window. Thermal images were taken at 1Hz before, during, and after illumination of the cortex. Regions of interest were drawn over the thermal images, and average temperatures calculated.

### Rose Bengal spectroscopy

To examine RB’s absorption spectra, we dissolved RB in 3 different solutions. In one sample, RB (Sigma Aldrich – 198250-5G) was dissolved (0.1mg/mL) in PBS (Cold Spring Harbor Laboratory Press) and its absorption spectrum was derived by a well plate reader (BioTek – Synergy H1) in increments of 1nm from 230nm-998nm. In another sample, RB was dissolved (0.1mg/mL) in PBS mixed with bovine serum albumin (VWR Life Sciences – CAS: 9048-46-8; 0.6mM concentration;Alarcón et al., 2009). To generate a plasma solution, mice to-be-culled were exsanguinated and their blood was mixed with heparin (Sandoz – DIN 02303108). The blood was centrifuged (Eppendorf – 5424) at 2000g for 10 minutes and the supernatant collected. RB was then dissolved in the plasma (0.1mg/mL) and the absorption spectrum was derived.

### Pre- and post-stroke vascular imaging

5 minutes before induction of photothrombotic stroke, maps of cortical blood flow were derived by laser doppler imaging (LDI) and laser speckle contrast imaging (LSCI) through the transcranial window. These maps were then also produced 5 minutes and 7 days after stroke induction.

### Photothrombotic stroke induction

A 10mg/mL Rose Bengal (RB) solution was prepared the day of stroke induction by dissolving RB (Sigma Aldrich – 198250-5G) in sterile PBS (Cold Spring Harbor Laboratory Press) within an opaque test tube. The solution was vortexed for 2 minutes and then sonicated for 10 minutes. Mice were anesthetized with isoflurane (5% induction and 2% maintenance in air) and head-fixed under a custom modular optical imaging system (LabeoTech – OiS200). A rectal temperature probe was inserted, and core temperature was maintained at 37±1.5°C via a feedback-regulated heating pad while the eyes were treated with eye lubricant (Optixcare) to keep the corneas moist. The light used for photothrombosis, either yellow (561nm @ 7mW/mm^2^ @ 2.2mm ⌀ (DPSS – MGL-U-561 – Changchun New Industries Optoelectronics Technology Co)) or green (520nm @ 7mW/mm^2^ @ 2.2mm ⌀ (DPSS – Labeotech Canada) was targeted 1.5mm lateral to bregma through the transcranial window via a system of galvo mirrors. RB was drawn up into a 31G insulin syringe (1633151B – Sol-M) about a minute before injection (100mg/kg I.P) into the prone mouse by lifting a hindleg. The RB was allowed to circulate for 3 minutes before the light was turned on for 13 minutes as previously reported (Schaffer et al., 2006; Harrison et al., 2013; Balbi et al., 2017).

### EEG monitoring

We recorded spreading depolarizations (SDs) by detection of the characteristic ultraslow direct-current (DC) shift, which results from differences in depolarization between soma and dendrites (Canals et al., 2005). The DC shift is both necessary and sufficient for identification of an SD (Dreier et al., 2017, 2019). Once mice were anesthetized, toothless copper alligator clips were fixed to the previously implanted stainless-steel electrodes, and silver wire (A-M Systems Cat No. 785500) was used to connect to the recording channel of a DC differential amplifier (A-M Systems Model 3000). A stainless-steel acupuncture needle (DongBang acupuncture – 0.25mm x 15mm) placed in the scruff of the mouse’s neck served as the reference electrode for the DC differential amplifier. Finally, another acupuncture needle placed in the mouse’s back served as a ground electrode and helped decrease noise. The DC-amplified signal (10kHz sampling rate, 60Hz notch filter, 0Hz high pass filter, 20kHz low pass filter, 100x gain) was digitized (LabeoTech – OiS200) and offline correction for signal drift and bandpass filtering (0.01-1Hz) was performed through the EEGLAB toolbox (Delorme and Makeig, 2004) for MATLAB (MathWorks).

In the optogenetic experiments, the EEG signal was amplified (A-M Systems Model 1700) with high pass filter set to 0.1Hz, low pass filter set to 20kHz, and the notch filter at 60Hz. Ten 20ms pulses of blue, green, or yellow light (450nm, 520nm, 561nm @ 7mW/mm^2^ @ 2.2mm ⌀, interpulse interval of 980ms) were delivered through the transcranial window in ChR2-expressing mice as well as non-opsin-expressing mice. Evoked EEG responses were averaged across mice but not processed any further. Control experiments were performed to ensure there was no photoelectric effect from the photothrombotic light (520nm or 561nm) on the EEG electrode (Fig.S1).

### Euthanasia, histology, and lesion volume

Histology was performed 7 days post-stroke. Mice were deeply anesthetized with 65mg/mL sodium pentobarbital (Bimeda-MTC – 00141704) and transcardially perfused with PBS (7 ml/min for 6min) and 10% neutral buffered formalin (7 ml/min for 10 min). Brains were extracted and fixed in formalin before serial sectioning on a vibratome (Vibratome – Series 1000 sectioning system) at 100µm and mounting on gelatin-coated slides. Slides were scanned by flatbed scanner (Canon – 9000F MKII) at a resolution of 1200dpi and also imaged on a microscope (Zeiss – ImagerM2) at 2x magnification for autofluorescence in the GFP channel. At this early survival time, the stroke was highly fluorescent and was manually traced in each section. Total lesion volume was calculated by summing the stroke volume from each section (stroke area x 100µm).

### Statistical analysis

All average traces were plotted using base R (version 4.4.2), where individual mice were normalized to a baseline recording period, and then averaged across animals. Sections from the normalized individual traces were averaged together and compared statistically as well as plotted using GraphPad Prism (version 10.6.0).

## Results

### Rose Bengal reduces cortical blood flow and cerebral blood volume

There are no reports of RB affecting cerebral blood flow in the absence of its photoactivation. However, any potential influence on the cerebral vasculature may be an intrinsic confound for the photothrombosis model. Thus, we monitored cortical blood flow through a transcranial window by LDF in isoflurane-anesthetized mice during administration of RB (Fig.1A left inset). First, an I.P injection of PBS was delivered as a control, followed 5 minutes later by an I.P injection of RB (opposite sides of the mouse). The initial PBS injection did not significantly affect cortical blood flow (Repeated measures F(1.633, 13.06) = 57.21, p < 0.0001, Dunnett’s p = 0.0725) while the subsequent RB injection resulted in a significant drop (Dunnett’s p < 0.0001; Fig.1A and right inset) that lasted on average at least 15 minutes. To complement our single point optical measurement of blood flow in the cortex and to also quantify potential hemodynamic changes in subcortical brain regions, we performed brain-wide measurements of perfusion using the doppler signal from functional ultrasound imaging (fUS; Fig.1B left inset and Movie 1). After RB we observed a significant drop in blood volume both cortically (Fig.1B; Repeated measures F(1.069, 5.345) = 36.77, p = 0.0013, Dunnett’s p = 0.0022) and subcortically (Fig.1C; Repeated measures F(1.053, 5.265) = 16.25, p = 0.0087, Dunnett’s p = 0.0158), whereas PBS elicited no change in cortical (Dunnett’s p = 0.7639) or subcortical (Dunnett’s p = 0.8178) blood volume. These data indicate that RB has intrinsic effects on blood flow/volume throughout the entire brain in anesthetized mice, independent of photoactivation.

**Figure 1.**
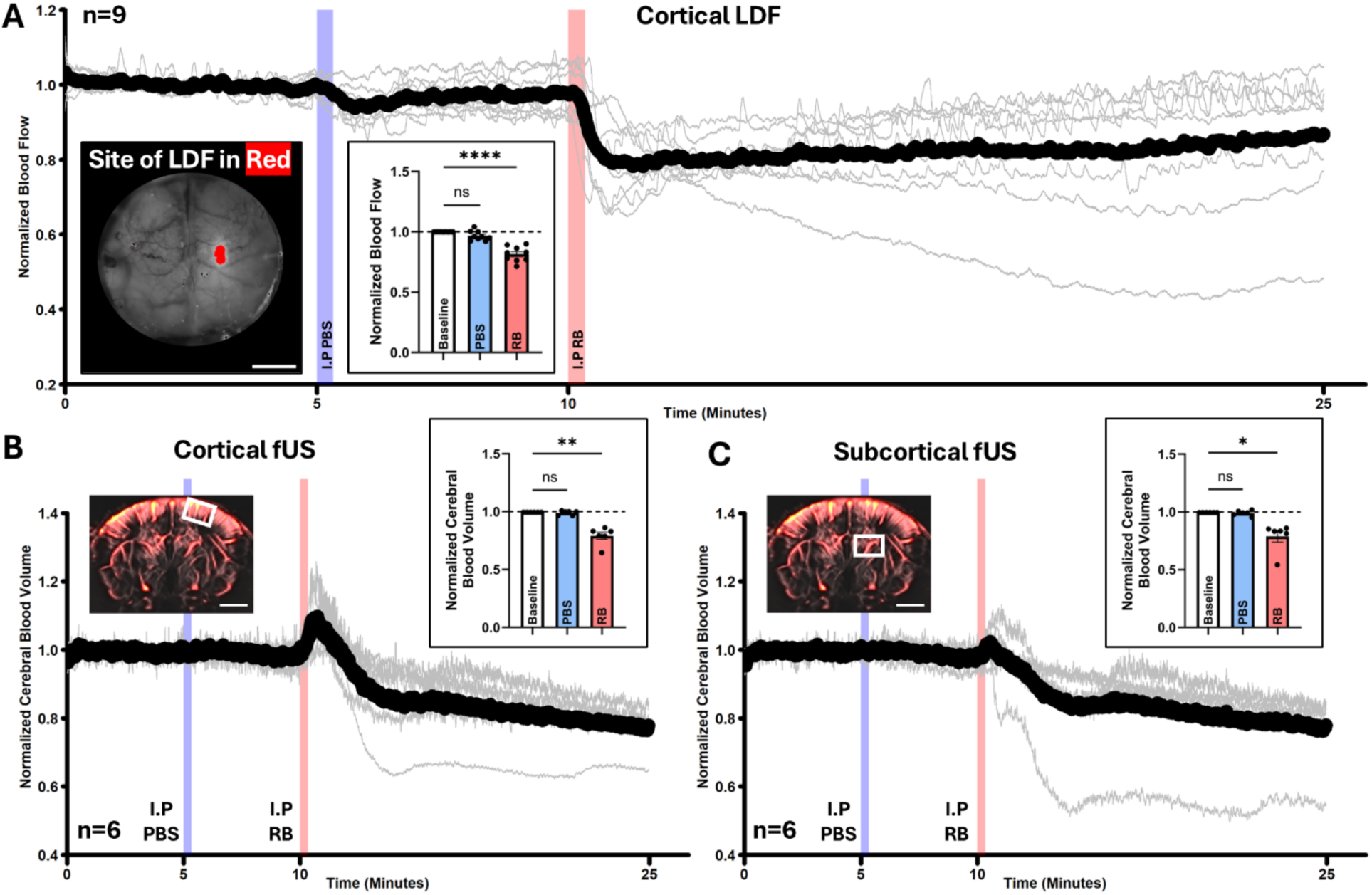
Rose Bengal reduces cortical blood flow and cerebral blood volume in the absence of photoactivation. **A**. Longitudinal LDF recording during baseline and following an I.P PBS (blue vertical shading) and RB injection (red vertical shading). Representative example of the recording site is visible in the left inset and quantification of the average flow is presented in the right inset (Baseline = 0:00 to 5:00, PBS = 5:00 to 10:00, and RB = 10:00 to 15:00). **B**. Cortical fUS of an I.P PBS injection (blue vertical shading) followed by an I.P RB injection (red vertical shading). Representative example of the ROI is visible in the left inset and quantification of the average blood volume is presented in the right inset (Baseline = 0:00 to 5:00, PBS = 5:00 to 10:00, and RB = 10:00 to 15:00). **C**. Subcortical fUS of an I.P PBS injection (blue vertical shading) followed by an I.P RB injection (red vertical shading). Representative example of the ROI is visible in the left inset and quantification of the average blood volume is presented in the right inset (Baseline = 0:00 to 5:00, PBS = 5:00 to 10:00, and RB = 10:00 to 15:00). Scale bars are 2mm, and the black trace in A, B and C represents the mean of the individual cases (grey traces).

### Rose Bengal constricts cortical blood vessels

The observations of RB diminishing blood flow and blood volume are surprising and suggest that RB may have intrinsic vasoactive properties. Indeed, a drop in cortical blood flow is typically associated with a preceding vasoconstriction (Devor et al., 2008; Hill et al., 2015; Hartmann et al., 2021), suggesting that RB may be influencing vascular tone. To verify this, we prepared mice with craniotomy windows and performed *in vivo* 2-photon microscopy under isoflurane anesthesia to directly measure changes in vessel diameter. We labeled the vasculature with retro-orbitally injected FITC-dextran (70kDa) and delivered either PBS or RB, all while imaging cortical vessels at 950nm (Fig.2A and B).

**Figure 2.**
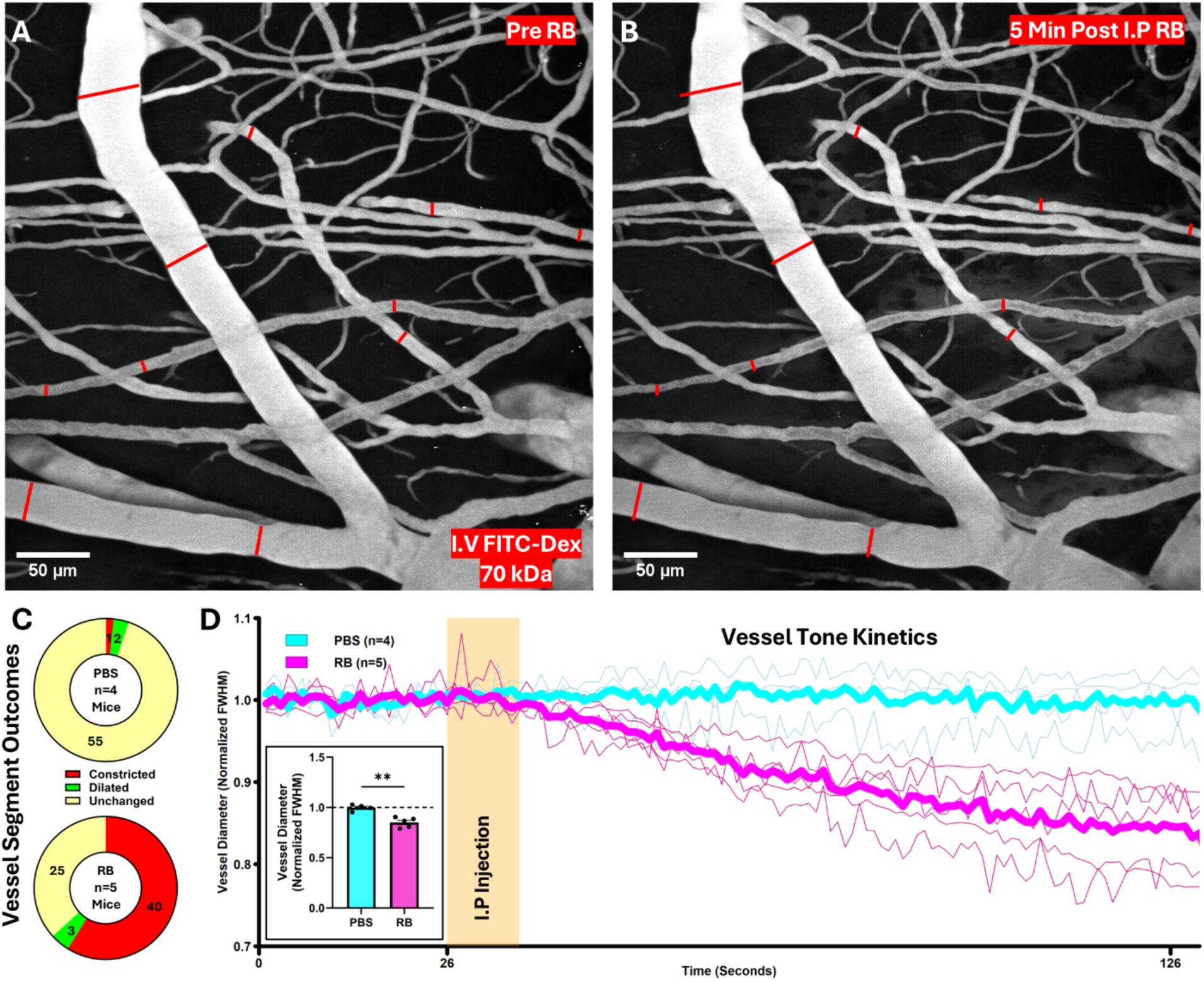
Rose Bengal constricts cortical blood vessels in the absence of photoactivation. **A**. Representative example of a maximum intensity Z-projection of FITC-Dextran labelled cortical blood vessels before RB injection. **B**. A Z-projection of the same cortical blood vessels taken 5 minutes after injection of RB. The red cross marks are the same length and in the same place in both images. **C**. A single vessel rich Z-plane was chosen and monitored throughout the injections, the number of vessels segments and their outcomes (constricted, dilated, and unchanged) across mice are shown after PBS (upper) and RB (lower) injections. **D**. Vessel diameter was monitored throughout PBS and RB injections and the average full width at half maximum (FWHM) for 3 randomly chosen vessel segments was calculated for each mouse (thin traces) and averaged together (thick traces). The inset compares the normalized FWHM averaged over the last 30 seconds of the recording for both injection groups.

To quantify vessel diameter, a vessel-rich Z plane was chosen and continuously recorded before, during, and after PBS or RB injections (Movie 2). By measuring the diameter of vessel segments throughout the injection, we classified vessel outcomes into three categories (constricted, dilated, or unchanged) and show that PBS causes no appreciable change in vessel diameter (Fig.2C upper) while RB causes the majority of vessels (59%) to constrict (Fig.2C lower). The kinetics of this RB-induced vasoconstriction (Fig.2D) indicate that cortical vessels begin to constrict within seconds of the I.P injection, and vessel diameter decreases to ∼85% of baseline 100 seconds after the injection (Fig.2D inset; Unpaired Welch’s t = 5.315, df = 6.796, p = 0.0012). Altogether, these data indicate that RB acts as a vasoconstrictor, even in the absence of photoactivation.

### Rose Bengal reduces peripheral blood flow

To determine if the observed hypoemia is specific to the brain, we applied our non-invasive measures of blood flow (LSCI, LDF, and LDI) to the superficial vessels of the hind paw in isoflurane-anesthetized mice (Fig.3A; Movie 3). Single point measurements made by LDF revealed that RB injection also causes a significant drop in peripheral blood flow (Fig.3B), whereas PBS injection does not (Fig.3B inset; Unpaired Welch’s t = 5.386, df = 6.437, p = 0.0013). Maps of blood flow acquired with LDI both before (Fig.3C and E) and after (Fig.3D and F) injections confirmed that there is a drop in flow across the entire paw after RB but not PBS (Fig.3G; Unpaired Welch’s t = 5.310, df = 7.154, p = 0.0010), indicating that the RB-induced drop in blood flow occurs in both the brain and the periphery.

**Figure 3.**
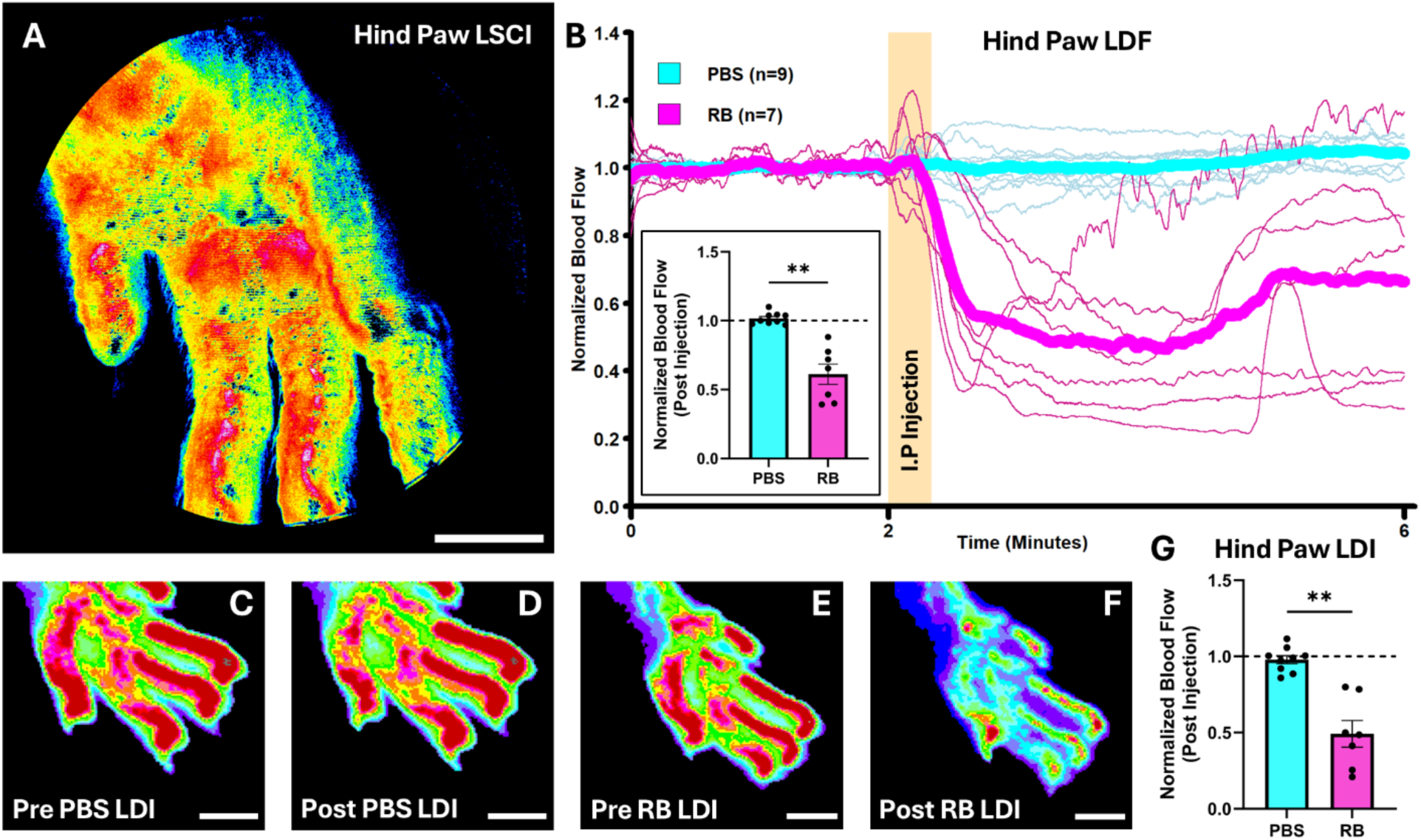
Rose Bengal reduces peripheral blood flow in the absence of photoactivation. **A**. Example of superficial vessels and blood flow in the mouse hind paw as measured by LSCI. **B**. Hind paw LDF during either PBS (cyan) or RB (magenta) injections. Thin traces show individual cases and thick traces show group averages. The LDF laser was positioned over the palm of the hind paw, and I.P injections were delivered to the opposite side of the mouse. The post injection period was average for each mouse and normalized to the average baseline flow (inset). **C**. Hind paw LDI taken before PBS. **D**. Hind paw LDI taken 5 minutes after PBS. **E**. Hind paw LDI taken before RB. **F**. Hind paw LDI taken 5 minutes after RB. **G**. LDI ROIs were traced over the entire hind paw, then an average value of flow was calculated. Post PBS and RB LDI paw averages were normalized to the baseline LDI paw averages for each mouse. Scale bars are 2mm.

### The Rose Bengal absorption spectrum

Previous reports suggest that RB exhibits a red-shifted (bathochromic) absorption spectrum when bound to albumin (Alarcón et al., 2009; Yoguim et al., 2022) or dissolved in blood plasma (likely bound to the albumin in the plasma) (Boquillon et al., 1992). Our spectrographic measurements confirm a redshift when RB was dissolved in bovine serum albumin (Fig.4;550nm to 561nm) or heparinized mouse plasma (550nm to 564nm). Thus, our results indicate that 561-564nm light (yellow) is optimal to activate intravascular RB.

**Figure 4.**
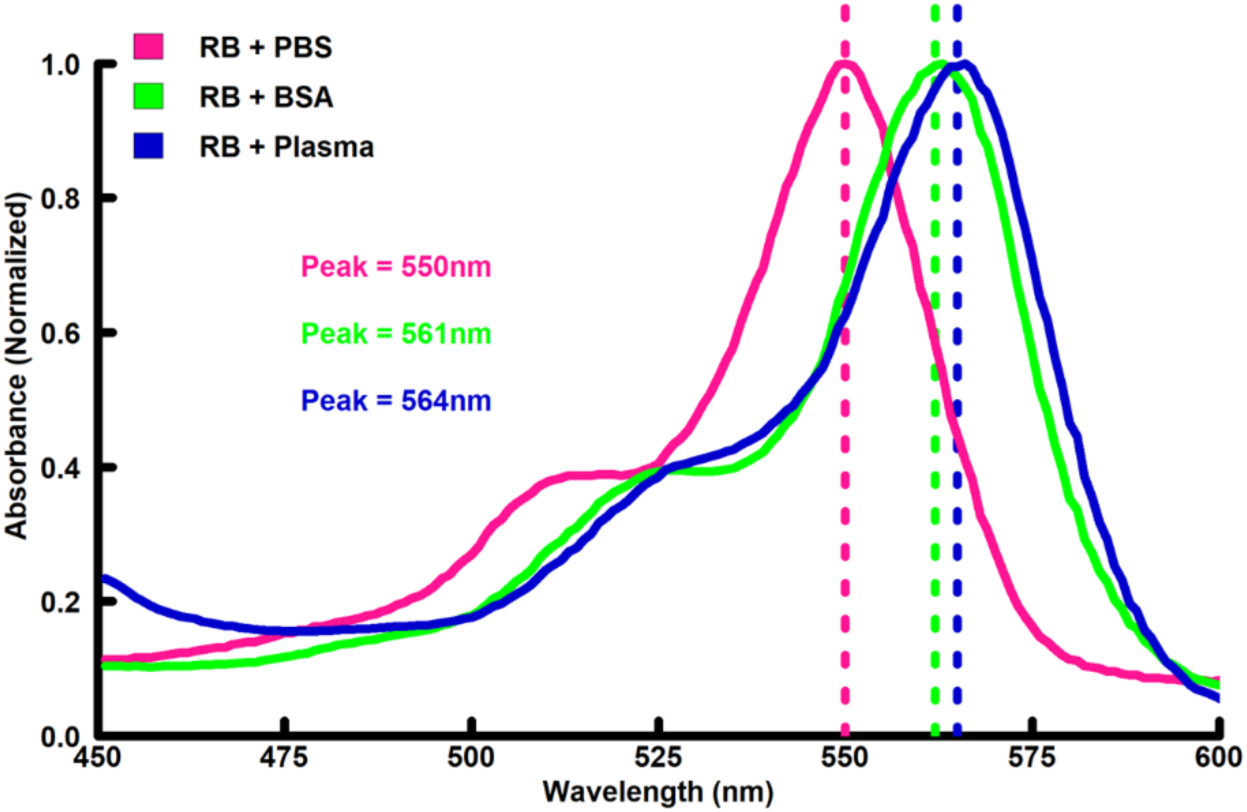
The RB absorption spectrum when dissolved in PBS, BSA, or blood plasma. BSA and blood plasma both caused a redshift in the absorption spectrum of over 10nm, making 561-564nm optimal for photoactivating RB *in vivo*.

### The light used for photothrombosis increases blood flow and brain temperature

During photothrombotic stroke, light is typically delivered to a fixed region at a 100% duty cycle for over 10 minutes, therefore making photothrombosis one of the most intensive applications of light during *in vivo* experiments. Although some stroke studies include light-only control groups (Kuroiwa et al., 2009; Clark et al., 2019; Knezic et al., 2022), the direct effect of photothrombotic light remains uncharacterized in this model. Thus, we monitored blood flow (LDF and LSCI) during 561nm illumination in the absence of any RB (with a typical photothrombosis timeline of 13 minutes of 7mW/mm^2^ light and a 2.2mm beam size). Our LDF recording showed a significant increase in blood flow at the site of illumination (Fig.5A; Paired t = 4.463, df = 13, p = 0.0006). To examine the areal extent of this blood flow increase, we performed LSCI and found a significant increase both at the site of illumination (Fig.5B; Paired t = 7.443, df = 13, p < 0.0001), as well as at the homotopic site in the opposite hemisphere (Fig.5C; Paired t = 9.044, df = 13, p < 0.0001). The exponential changes in flow were coincident with the onset and offset of the laser and were reminiscent of previously reported light-induced temperature changes in the brain (Christie et al., 2013; Stujenske et al., 2015; Arias-Gil et al., 2016; Owen et al., 2019). Thus, we decided to image temperature through the transcranial window during continuous illumination (Fig.5D). We found that brain temperature begins to rise immediately when the light is turned on and continues to rise by ∼3°C within 30 seconds relative to baseline (Paired t = 23.99, df = 5, p < 0.0001). Surprisingly, even in the homotopic region of the contralateral hemisphere, a temperature increase of ∼1.5°C was observed (Paired t = 7.805, df = 5, p = 0.0011). Altogether, these data suggest that the light used for photothrombosis introduces hyperemia and tissue heating.

**Figure 5.**
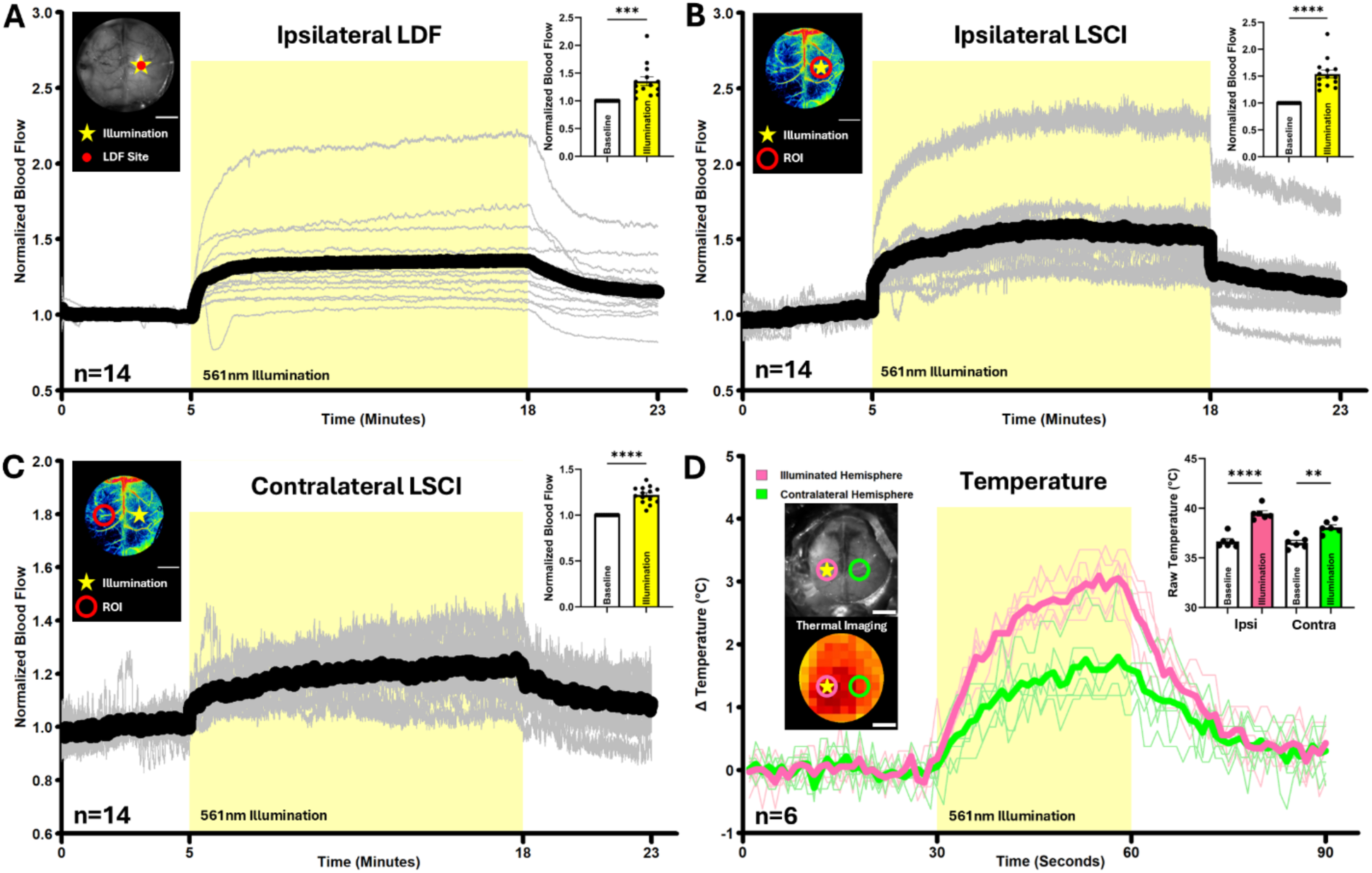
Photothrombotic light illumination results in hyperemia and tissue heating. **A**. Ipsilateral transcranial LDF of blood flow during 561nm light illumination. The infrared laser was positioned in the centre of the 561nm illumination site. The average of baseline (0:00-5:00) was compared to the average at steady state (13:00-18:00). **B**. Ipsilateral transcranial LSCI of blood flow during 561nm light illumination. An ROI was drawn over the illuminated site. The average of baseline (0:00-5:00) was compared to the average at steady state (13:00-18:00). **C**. Contralateral transcranial LSCI of blood flow during 561nm light illumination. An ROI was drawn over the region of the opposite hemisphere homotopic to the light stimulation site. **D**. Transcranial thermal imaging of both hemispheres during 561nm illumination. An ROI was drawn over the illuminated site (pink) and the region of the opposite hemisphere homotopic to the illumination site (green). All mice had a baseline brain temperature of 37±1.5°C. The average of baseline (0:00-0:30) was compared to the average temperature just before the light was turned off (00:45-00:60). Scale bars are 2mm.

### The magnitude of blood flow drop, lesion volume, and the onset of spreading depolarizations are distinctly impacted by yellow and green light during photothrombosis

Isoflurane-anesthetized mice with transcranial chronic windows received a photothrombotic stroke while monitoring hemispheric DC-EEG and blood flow in the core of the stroke via LDF. Strokes were produced by either green (520nm) or yellow light (561nm), while ensuring that laser intensity (7mW/mm^2^), beam size (2.2mm), and duration of illumination (13 min) were the same for both groups.

While blood flow was stable at baseline for both the green (Fig.6A) and yellow (Fig.6B) photothrombosis groups, injection of RB (at 2 minutes) caused a visible drop in flow which persisted until the time of illumination. This decrease in flow preceded any photoexcitation and was observed in both green and yellow stroke groups (Fig 6C; Unpaired Welch’s t = 1.700, df = 30.37, p = 0.0993). Upon illumination with green or yellow light (at 5 minutes), flow initially rose before beginning to drop over the remaining illumination time. The initial rise in flow was statistically identical between green and yellow strokes (Fig.6D; Unpaired Welch’s t = 1.095, df = 31.95, p = 0.2816). However, yellow light produced a significantly larger decrease in blood flow than green light (Fig.6E; Unpaired Welch’s t = 6.941, df = 20.11, p < 0.0001). To complement our LDF measures, blood flow was also recorded with LSCI in another cohort of mice (n=4; Movie 4) and results showed a similar hemodynamic profile when measured at the site of the stroke. Lesion volume measurements 7 days post-stroke showed that green light strokes were significantly smaller than yellow light strokes (Fig.6F; Unpaired Welch’s t = 5.398, df = 31.92, p < 0.0001).

**Figure 6.**
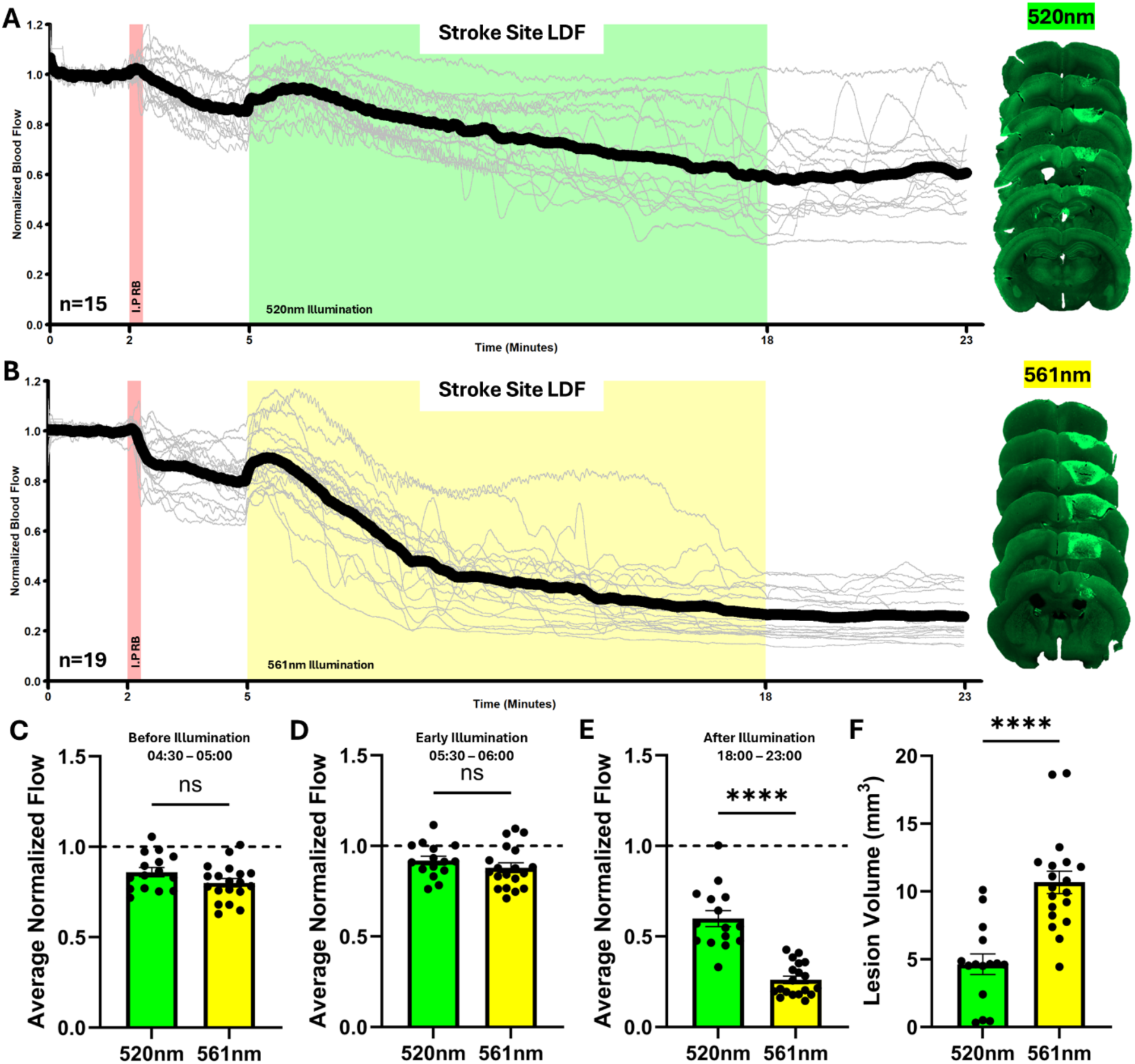
The hemodynamic progression of green (520nm) and yellow (561nm) light photothrombotic strokes. **A**. Normalized LDF recorded through the transcranial window during green light photothrombosis. The infrared laser was targeted within the centre of the photothrombotic laser, giving a measurement of cortical blood flow in the core of the stroke. The red vertical shading indicates the injection of RB. The green shading indicates the time for which the photothrombosis laser was on (520nm @ 7mW/mm^2^ @ 2.2mm ⌀). On the right is a representative brain collected 7 days after the stroke (300µm apart). **B**. Normalized LDF recorded through the transcranial window during yellow light photothrombosis (561nm @ 7mW/mm^2^ @ 2.2mm ⌀). For A and B thin traces show individual cases and thick traces show group averages. **C**. Normalized flow averaged over the last 30 seconds before green and yellow light illumination. **D**. Normalized flow averaged from 5:30-6:00 minutes. **E**. Normalized flow averaged over the entire post illumination period. **F**. Lesion volume measured 7 days after stroke for both green and yellow light strokes.

The electrophysiological progression of the green and yellow light strokes was monitored by epidural DC-EEG. The occurrence of spreading depolarizations (SDs) was manually quantified as any event that caused a deflection in the EEG trace greater than 0.5mV. During photothrombosis with yellow light, more SDs propagated (Fig.7C; Unpaired Welch’s t = 2.212, df = 22.89, p = 0.0372) and began sooner after illumination (Fig.7D; Unpaired Welch’s t = 6.291, df = 15.98, p < 0.0001) than with green light. However, there was no difference between groups in the frequency of SDs (Fig.7E; Unpaired Welch’s t = 0.5588, df = 22.46, p = 0.5819), or the level of ischemia at the first SD event (Fig.7F; Unpaired Welch’s t = 1.932, df = 23.96, p = 0.0653).

**Figure 7.**
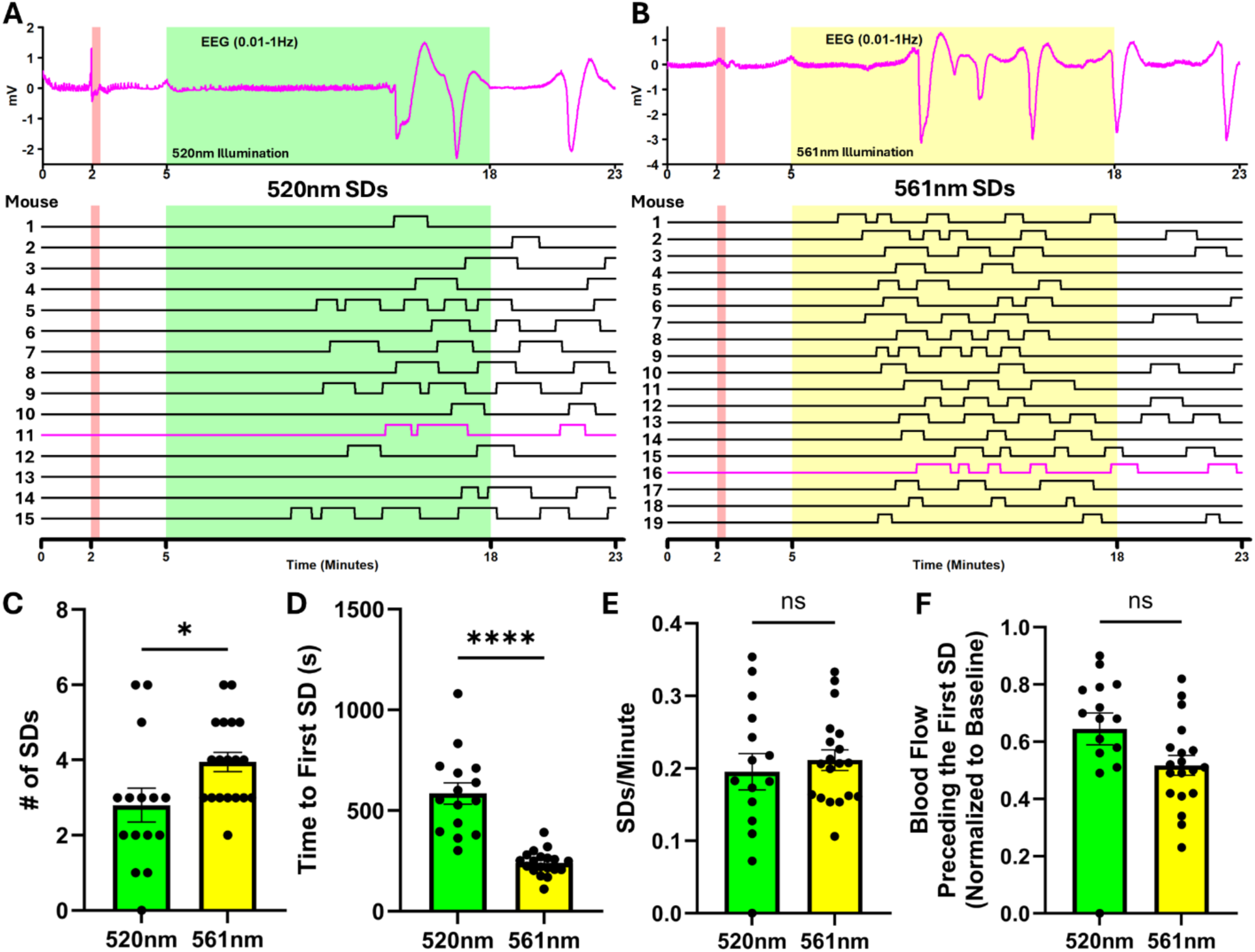
The DC-EEG progression of green (520nm) and yellow (561nm) light photothrombotic strokes. **A**. Filtered (0.01-1Hz) EEG during green light photothrombosis (upper) (red vertical shading indicates the I.P RB injection and green shading indicates the period of light illumination). The DC shifts (presumed to be individual SDs) present in the EEG traces are reported for each mouse below (upper EEG trace is mouse #11), where upticks represent the period of each SD. **B**. Filtered (0.01-1Hz) EEG during yellow light photothrombosis (upper). The DC shifts (presumed to be individual SDs) present in the EEG traces are reported for each mouse below (upper EEG trace is mouse #16), where upticks represent the period of each SD. **C**. The total number of SDs recorded during green and yellow light photothrombosis induction. **D**. The time delay from green or yellow light illumination to the first SD. **E**. The frequency of SDs calculated from the first one to the end of the recording for each mouse. **F**. The normalized blood flow value (LDF) at which the first SD occurred.

### Green light alters blood flow and the frequency of spreading depolarizations during photothrombosis in ChR2 expressing-mice

Owing to the significant advantages offered by all-optical techniques, photothrombotic strokes are often induced in opsin-expressing mice (Harrison et al., 2013; Paz et al., 2013; Anenberg et al., 2014; Lim et al., 2014; Shah et al., 2017; Tennant et al., 2017; Wahl et al., 2017; Allegra Mascaro et al., 2019; Bo et al., 2019, 2020; Zhang et al., 2019, 2021; Abe et al., 2021; Balbi et al., 2021; Lin et al., 2021; Mirza Agha et al., 2021; Bice et al., 2022; Conti et al., 2022; Suo et al., 2023; Wang et al., 2023; Okabe et al., 2025). One unreported confound is any potential crosstalk between the light used for stroke induction (typically green) and photoactivation of the opsin (typically ChR2). It is unclear however, whether green light (520nm), when used at a photothrombosis appropriate intensity, is capable of driving ChR2. We began by attempting to optically evoke EEG deflections in ChR2-expressing mice (Thy1-ChR2-YFP; Fig.8A) using either blue (450 nm), green (520 nm), or yellow (561nm) light pulses (20ms duration, 980ms interpulse interval; @ 7mW/mm^2^). As expected, (Boyden et al., 2005; Zhang et al., 2006; Lin, 2011; Mattis et al., 2012; Pan et al., 2014), blue light produced the largest deflections (Fig.8A upper). However, green light also produced a slightly smaller deflection with 100% fidelity to light pulses (Fig.8A middle), while yellow light did not produce detectable deflections (Fig.8A lower). Pulsing experiments were also repeated in opsin-negative mice, where no deflections were observed under any condition (Fig. S2).

**Figure 8.**
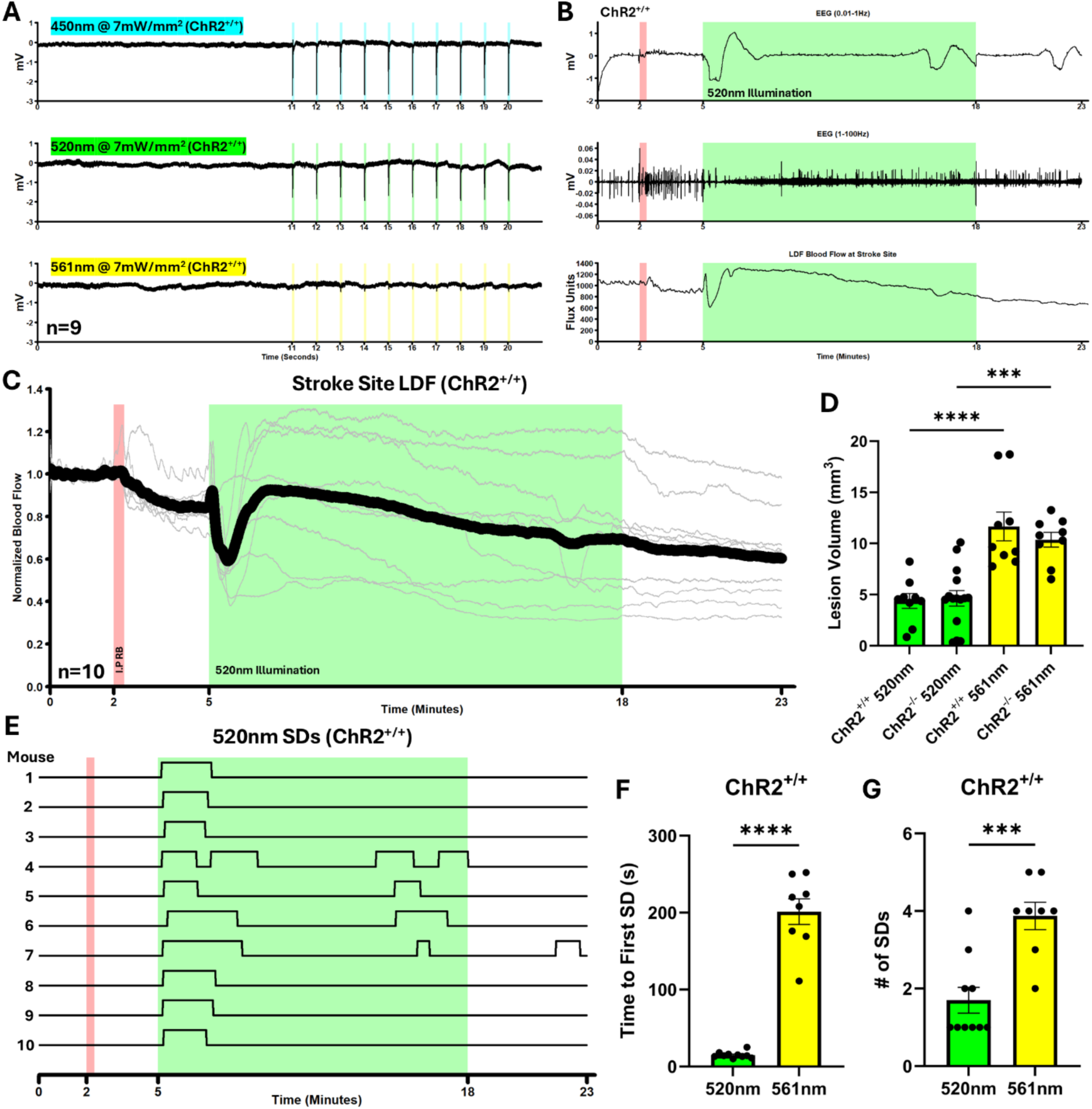
The Crosstalk between photothrombosis and ChR2. **A**. Blue (450nm), green (520nm), and yellow (561nm) light-evoked EEG deflections in ChR2 expressing mice. 20ms (interpulse interval of 980ms) light pulses were delivered and the traces from 9 mice were averaged together. **B**. Representative example of aligned EEG and LDF traces from one ChR2 expressing mouse during green light (520nm) photothrombosis. EEG is filtered into its low (DC shifts) and high frequency (general cortical activity) components. **C**. Normalized LDF recorded through the transcranial window during green light (520nm) photothrombosis in ChR2 expressing mice. The infrared laser was targeted within the centre of the photothrombotic laser, giving a measurement of cortical blood flow in the core of the stroke. The red vertical shading indicates the I.P injection of RB. The green shading indicates the time for which the photothrombosis laser was on (520nm @ 7mW/mm^2^ @ 2.2mm ⌀). **D**. Lesion volumes from ChR2 expressing and control mice (both green (520nm) and yellow (561nm) light) collected 7 days after stroke. **E**. The occurrence of SDs across all ChR2 expressing mice. Each SD corresponds to an individual DC shift from the EEG. **F**. Time delay from light onset to first SD for both wavelengths of light in ChR2 expressing mice. **G**. The number of SDs (DC shifts) recorded by EEG throughout the intraoperative monitoring period for both wavelengths in ChR2 expressing mice. Of note is that there were no differences for any outcome between 561nm ChR2 expressing and ChR2 non expressing mice.

To directly examine the effect of ChR2 activation during photothrombosis on stroke outcome, we performed green light and yellow light photothrombosis in ChR2-expressing and non-opsin-expressing mice. Blood flow at the stroke site (LDF) as well as hemispheric DC-EEG (representative aligned traces in Fig.8B) were recorded continuously. Blood flow in ChR2 expressing mice showed the same initial pattern as in non-opsin expressing mice (Fig 6a), including a decrease immediately after RB injection. However, within ∼10s of green light illumination, a large ischemic transient (Fig.8B lower), a downward shift of the DC potential (Fig.8B upper) or depression of the spontaneous EEG occurred in 100% of the mice (Fig.8B middle). This likely indicates the initiation of an optogenetic SD event, the propagation of which was confirmed by LSCI (Movie 5). The hemodynamic profile of this optogenetic SD was consistent across mice (Fig.8C), while the ensuing drop in blood flow varied across subjects. Of note is that despite the addition of an optogenetic SD at the beginning of the stroke induction, no difference in lesion volume was found between ChR2-expressing and non opsin-expressing mice (Fig.8D; F (3, 38) = 16.29, p < 0.0001, Tukey’s p = 0.9973), while illuminating with yellow light (561nm) resulted in a lesion volume increase over the green light for both the opsin-expressing (Tukey’s p < 0.0001) and non-expressing mice (Tukey’s p = 0.0003). Similarly, the SD profile was also dependent on the wavelength of PT light (Fig.8E). Although green light activation of ChR2 produced an SD nearly immediately after illumination began (Fig.8F; Unpaired Welch’s t = 11.10, df = 7.089, p < 0.0001), this opto-SD often seemed to “block” the initiation of further spontaneous SDs, resulting in significantly fewer SDs relative to yellow light (Fig.8G; Unpaired Welch’s t = 4.487, df = 15.54, p = 0.0004). The optogenetic SD was absent and no unexpected electrophysiological or hemodynamic confounds were seen when strokes were induced with yellow light (561nm) in ChR2-expressing mice (Fig.S2).

### Characterizing sex differences in lesion volume, longitudinal blood flow changes across both hemispheres and the occurrence of SDs following yellow light photothrombosis

To further examine a trend of male mice having larger strokes than female mice (Fig.S3A and B), we performed an additional experiment that also quantified SDs during photothrombosis and blood flow changes 7 days later (Fig.S4). Lesion volume measurements showed that male strokes were significantly larger than female strokes (Fig.9A inset and right; Unpaired Welch’s t = 3.011, df = 18.74, p = 0.0073). We pooled this cohort with the two previous green and yellow light stroke cohorts, and show that across experiments, males have significantly larger strokes (Fig.S3C).

**Figure 9.**
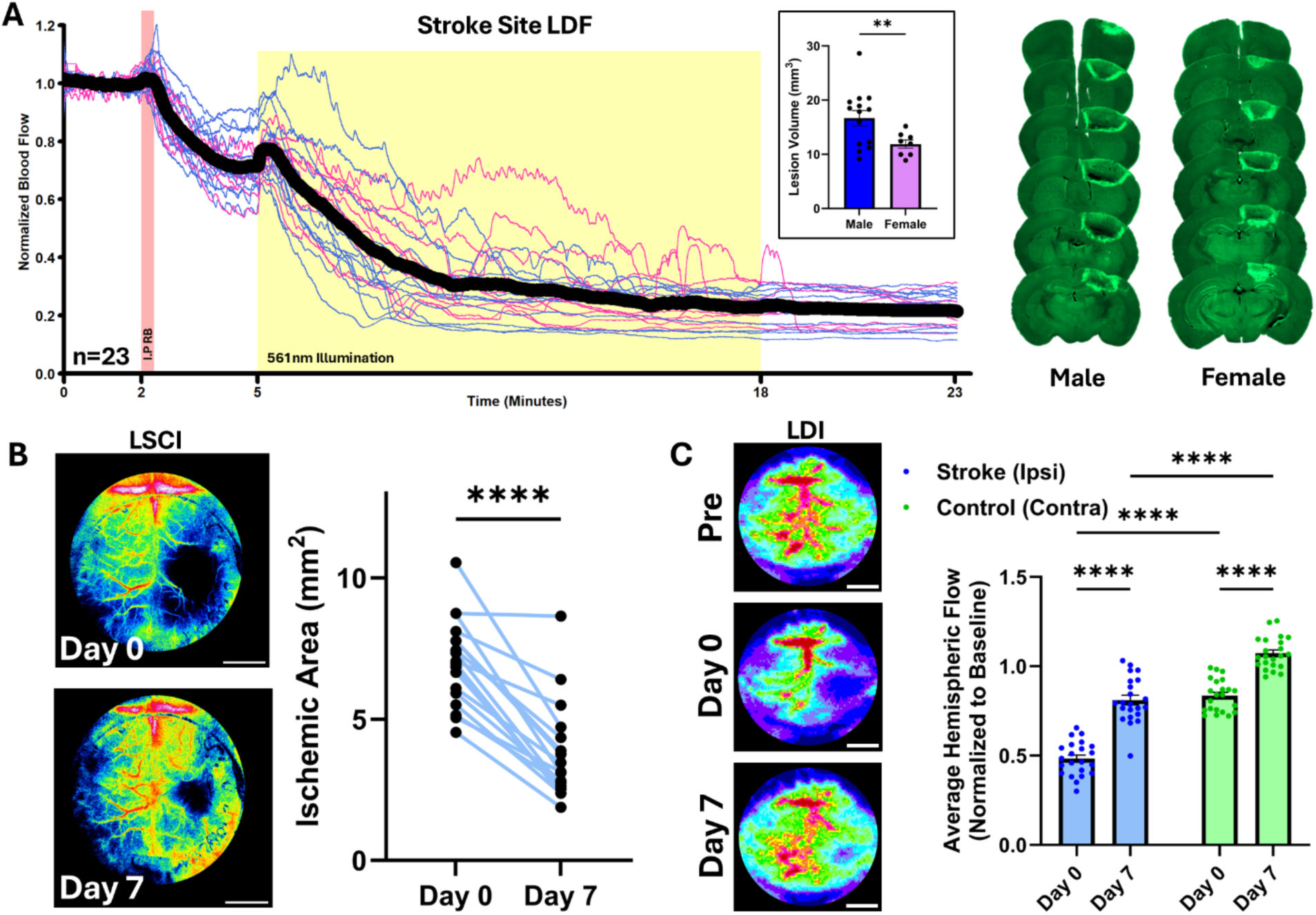
Hemodynamic sex differences and lesion progression in photothrombosis. **A**. Normalized LDF recorded through the transcranial window during yellow light photothrombosis. The infrared laser was targeted within the centre of the photothrombotic laser, giving a measurement of cortical blood flow in the core of the stroke. The red vertical shading indicates the injection of RB where a hind paw was lifted and an I.P injection was made. The green shading indicates the time for which the photothrombosis laser was on (561nm @ 7mW/mm^2^ @ 2.2mm ⌀). Thin pink traces are the female mice, and thin blue traces are the male mice (no difference between sexes) while the inset shows the lesion volumes between the sexes. On the right are representative examples of male and female lesions (300µm apart). **B**. The area of ischemia (right) measured by manual tracing of LSCIs made immediately after stroke induction and 7 days later (left). **C**. LDIs were taken just before, just after, and 7 days after stroke induction (left). ROIs were drawn over either the right or left hemisphere, and an average value of hemispheric blood flow was calculated and normalized to the pre stroke value. On the right, the average normalized hemispheric flow is presented. Scale bars are 2mm.

**Figure 10.**
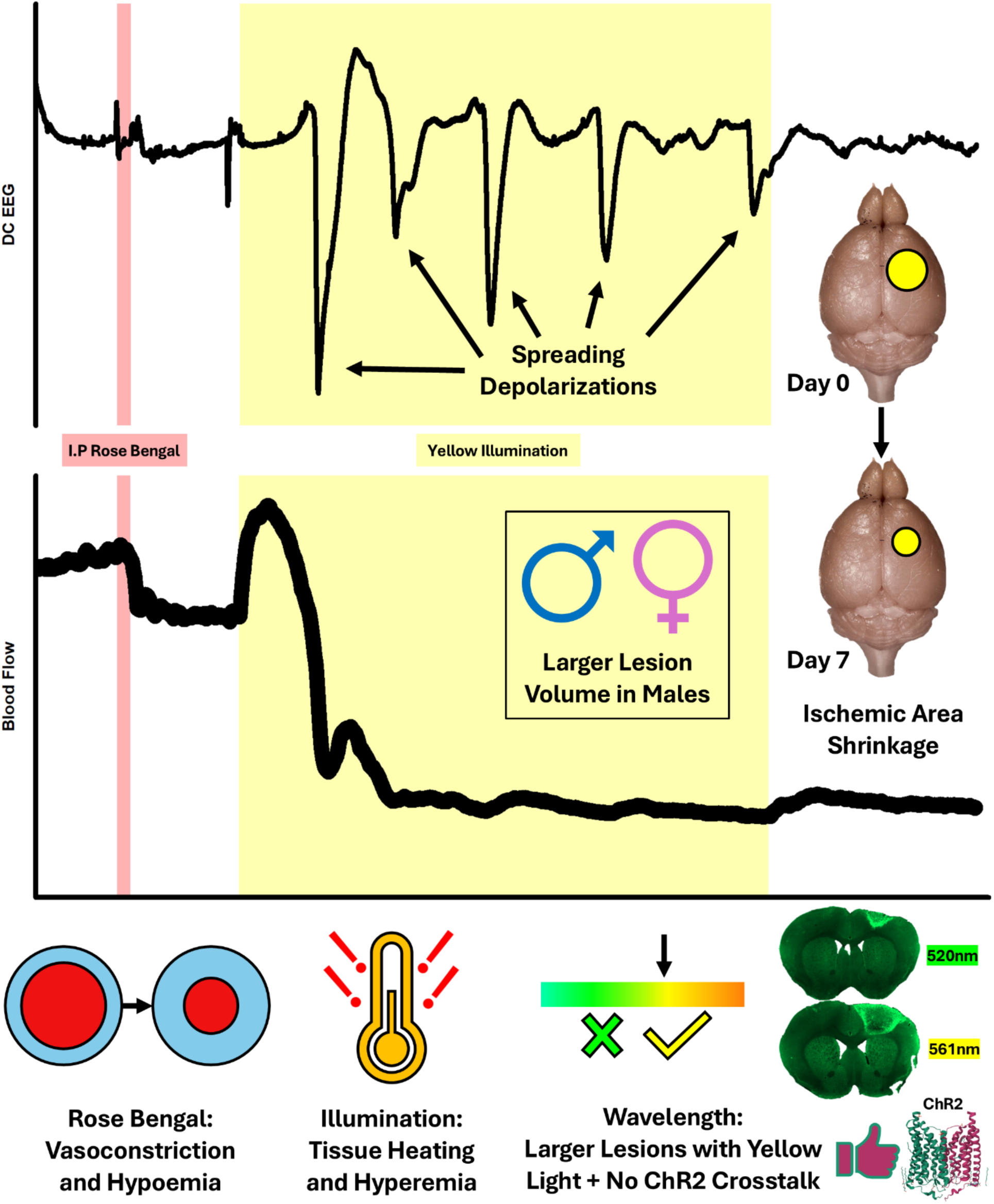
Cartoon depicting the hemodynamic and electrophysiological progression of the Rose Bengal photothrombotic stroke in mice, indicating potential caveats and confounds such as RB induced vasoconstriction, tissue heating and hyperemia, larger strokes with yellow light, wavelength crosstalk with ChR2, lesion maturation, as well as sex differences in lesion volume.

The area of ischemia was determined by LSCI both immediately after photothrombosis, and 7 days later, just before sacrifice. Our data shows that the photothrombosis-induced ischemic area decreases in size by 7 days after induction (Fig.9B; Paired t = 9.676, df = 18, p < 0.0001), similar to previous studies (Ginsberg and Busto, 1989; Lee et al., 2007; Kuroiwa et al., 2009; Labat-gest and Tomasi, 2013; Li et al., 2014; Aswendt et al., 2021). Along with LSCI, average measurements of blood flow were recorded for both the ipsilesional and contralesional hemispheres using LDI (Fig.9C). The average blood flow in the stroke hemisphere dropped to ∼50% of baseline immediately after stroke induction and returned to ∼80% of baseline by 7 days post stroke. Conversely, in the contralesional hemisphere blood flow dropped to ∼80% of baseline immediately after stroke induction, before increasing to ∼105% of baseline by 7 days post stroke.

We next wanted to examine any potential relationship between the decrease in blood flow observed immediately after RB injection and ischemic area (LSCI at both day 0 and 7) as well as final lesion volume (Fig.S5). We found significant correlations between ischemic area (both at day 0 and 7) and magnitude of blood flow decrease after RB injection. Although it appears that a larger drop in blood flux is predictive of a larger ischemic area, it is important to note that this relationship is driven by cases of failed stroke induction where we observed a lesion less than 2mm^3^ and less than 10% drop in flow. These data suggest that blood flow needs to decrease beyond a threshold of around 15% after RB injection for successful photothrombosis. However, the absolute drop in flow beyond that threshold is not predictive of lesion size. Indeed, no other hemodynamic metric predicted the size of the stroke or our observed sex difference in lesion volume.

We also investigated the progression of SD waves as a predictor of stroke outcome. SDs (indicated by the signature DC shift) were frequent following illumination (Fig.S6A upper), and we also observed the concomitant depression of spontaneous EEG (Fig.S6A middle). Interestingly, the first DC shift coincides with a large ischemic transient measured by LDF (Fig.S6A upper and lower). However, any subsequent SDs elicited very little change in blood flow, likely because the ischemic core (where LDF is being measured) no longer had the capability of propagating the metabolically-demanding SD. Indeed, when LSCI measurements were performed at more distal sites within the same hemisphere (Movie 4), hemodynamic transients (often increases in blood flow) were coincident with all DC shifts and not only the first one. Separating the traces of SDs by sex (Fig.S6B) revealed no differences in number of events (Fig.S6C; Unpaired Welch’s t = 1.139, df = 20.79, p = 0.2678), onset delay to the first event (Fig.SD; Unpaired Welch’s t = 1.341, df = 20.30, p = 0.1947), frequency (Fig.S6E; Unpaired Welch’s t = 1.208, df = 21.71, p = 0.2401), or level of ischemia at the first SD (Fig.6SF; Unpaired Welch’s t = 0.4128, df = 19.03, p = 0.6844). Importantly, none of these metrics predicted lesion volume in either sex.

## Discussion

Our experiments characterize several previously under/unreported caveats and confounds related to RB-mediated photothrombosis. We found that I.P RB constricts blood vessels and induces a drop in blood flow across the brain and periphery (hind paws), while light stimulation alone significantly increases cortical blood flow and temperature. We demonstrate that the use of the optimal wavelength of light during photothrombosis (yellow-561nm) improves stroke consistency by producing a larger, more pronounced drop in blood flow, ultimately resulting in a larger stroke. Importantly, in experiments involving ChR2 expression, yellow light illumination also eliminates ChR2 overactivation, which we show occurs during green light illumination. Lastly, we note across stroke cohorts that photothrombosis produces a larger stroke in male mice than female mice.

### The vasoactive properties of Rose Bengal

Across all studies which monitor hemodynamics during photothrombosis, there are no previous reports of RB’s hemodynamic effect (Yao et al., 2003; Kuluz et al., 2007; Kuroiwa et al., 2009; Wang et al., 2010; Lu et al., 2014; Seto et al., 2014; Qiu et al., 2016; Ling et al., 2018; Sullender et al., 2018; Clark et al., 2019; Sun et al., 2020; Sunil et al., 2020; Hu et al., 2022; Raman-Nair et al., 2023). We suspect this is due to incomplete recordings in the previous studies. In some cases, baseline monitoring of blood flow began after the injection of RB, possibly masking the relatively quick drop in flow we have observed, and in other cases, the animals were removed from blood flow monitoring, injected with RB while supine, and then returned prone to the monitoring equipment (creating a gap in the recording). Similarly, studies investigating the 2-photon excitation of RB for the purpose of microvessel occlusion also omit any mention of RB’s vasoconstrictive properties (Fukuda et al., 2022; Delafontaine-Martel et al., 2023; Zhu et al., 2023a, 2023b).

The confounding effects of RB can be seen at many levels. For instance, the full body hypoemia may have neuroprotective effects via ischemic conditioning, a phenomenon which is well-documented in animals (Weir et al., 2021) and to some degree in humans (Blauenfeldt et al., 2024; Ganesh and Testai, 2024; Keevil et al., 2024). Although the mechanisms underlying the protective effects of ischemic conditioning are unclear, they may be an essential (and confounding) part of the photothrombotic model. Furthermore, even small and regional drops in cerebral blood flow are associated with altered brain physiology (Hauglund et al., 2025) and behaviour (Abe et al., 2021). Finally, a global drop in cerebral blood volume may cause global disruptions to the functional connectivity of the brain in addition to the disruptions happening due to the stroke itself (Bandet and Winship, 2024).

### Light-induced tissue heating and hyperemia

We found significant tissue heating and hyperemia upon illumination with the photothrombotic light alone. These effects were not merely localized to the site of stimulation but were measurable even in the opposite hemisphere, well beyond the stroke site. In the context of stroke, modest post-ischemic hyperthermia is known to worsen outcomes (Kim et al., 1996; Reglodi et al., 2000; Sinigaglia-Coimbra et al., 2002), as is intraoperative hyperthermia (Busto et al., 1987). More broadly, we are not the first to report light-induced hyperemia and brain heating (Christie et al., 2013; Allen et al., 2015; Stujenske et al., 2015; Arias-Gil et al., 2016; Rungta et al., 2017; Owen et al., 2019; Peixoto et al., 2020); however previous reports have focused on short-duration pulsed light (which is commonly used for optogenetics) as opposed to long-duration 100% duty cycle photothrombotic light. While the specific long-term consequences of such hyperemia and tissue heating are unclear, considering that nearly all fundamental electrophysiological properties are temperature-dependent, such as action potential waveforms (Hodgkin and Katz, 1949) and channel conductances (Hille, 2001), it is fair to say that heat is not inert (Podgorski and Ranganathan, 2016 for a review on this topic). It would be interesting to evaluate the impact of heat and hyperemia on sensitive measures of circuit function or behaviour, separate from stroke-induced deficits.

### The design of sham strokes

For photothrombosis studies, the sham group typically consists of one of the following: a completely naïve animal, an animal who receives a saline injection followed by light illumination, an animal who only receives light illumination, or an animal who receives RB without light illumination. We have shown that both individual manipulations of the photothrombotic stroke, namely the RB injection and the light illumination, have their own separate and non-negligible effects on the progression of the stroke. Of note is that the RB induced hypoemia and the light induced hyperemia work in opposite directions and may compete with one another when done in sequence as opposed to individually. However, to capture as much of these dynamics as possible, both manipulations should be part of any control procedure. Thus, the optimal photothrombotic sham is then an animal who receives light illumination followed by an injection of RB. Furthermore, a non-manipulated naïve control group would also be beneficial for most experimental designs.

### Stroke volume

One of the goals for better aligning preclinical and clinical stroke research is to have models which recapture the relative size and location of human stroke in animals (Corbett et al., 2017). One criticism of photothrombotic stroke is that they are often small and mostly affect the superficial layers of cortex. With yellow light, our strokes are very comparable in volume to those produced by other well-established models (e.g. the largely cortical distal middle cerebral artery occlusion (Llovera et al., 2014; Leng et al., 2023; Marks et al., 2025)), but with the added benefit of being highly targetable. Importantly, these larger photothrombotic lesions were produced by using previously-established and accepted experimental parameters (100mg/kg I.P RB and 13 minutes of 7mW/mm^2^ light through intact skull). Perhaps by increasing the concentration of RB, the duration of light, the intensity of light, or thinning the skull, similarly sized strokes could be achieved using green light; however, as our data suggests, each of these choices comes with significant outcome-relevant drawbacks. Altogether, the use of yellow light during photothrombosis is a simple way of making larger strokes without added confounds.

In our study, we show that male strokes are larger than female strokes. While we investigated potential intraoperative factors as a potential mechanism (e.g. magnitude of initial ischemia, number of SDs) these factors did not predict our observed difference (or the variability in lesion volume more generally). Perhaps some postoperative metric (for example total number of SDs experienced during recovery) would be a better predictor of final lesion volume. Our observed sex difference may result from the neuroprotective effects of estrogen (Behl et al., 1997; Simpkins et al., 2012; Tang et al., 2022) and may be an intrinsic feature of the photothrombotic model. Indeed, there are reported sex differences in human stroke outcomes, although the causes are unclear (Dula et al., 2020; Rexrode et al., 2022; Owais et al., 2024). Previous photothrombotic studies show either no difference in lesion volume (Newton et al., 2022; Raman-Nair et al., 2023) or smaller female strokes (Cai et al., 1998; White et al., 2025), in line with what we have seen. However, most studies do not stratify lesion volume by sex or only use animals of one sex (typically male), limiting our ability to compare results.

Previous studies have documented that within the photothrombotic model, the final lesion volume evolves over time (Ginsberg and Busto, 1989; Lee et al., 2007; Kuroiwa et al., 2009; Labat-gest and Tomasi, 2013; Li et al., 2014; Aswendt et al., 2021). In particular, the lesion increases in size immediately following induction (doubles in volume within a few hours), peaking at 24-48 hours, and steadily decreases in volume thereafter. This pattern of lesion progression is different from the middle cerebral artery occlusion model, where lesion size increases modestly in the hours following induction (Bardutzky et al., 2005; Henninger et al., 2006), and changes very little thereafter (Virley et al., 2000; Cotrina et al., 2017). This difference in stroke maturation between the two models may be an important factor in studies assessing neuroprotective interventions or recovery mechanisms.

Much great work has been done to understand the pathophysiology of the photothrombotic model as well as to improve upon its basic form. Through a complementary and thorough combination of optical, ultrasonic, microscopic, electrophysiological, and histological investigations, we have described several hitherto unreported caveats of the photothrombotic model. It will be important for future studies (as well as past findings) to account for the vasoactive properties of RB, light-only heating effects, wavelength used to excite RB (both with and without ChR2 present), and sex differences in final lesion volume when designing experiments and interpreting data. Our study provides a framework for investigating the progression of photothrombosis during the induction and immediate aftermath of injury. Extending these hemodynamic and electrophysiological recordings beyond the intraoperative period will be important for studies evaluating long-term recovery after photothrombosis. Altogether, our study significantly advances our understanding of the photothrombotic model, and how best to use it.

## Conflict of interest statement

The authors declare no competing financial interests

## Acknowledgements

This work was supported by a Canadian Institute for Health Research project grant and Canadian Foundation for Innovation grant to G.S; uOttawa Brain and Mind Research Institute scholarship to P.C. We thank Zachary Eckert for his stroke pilot studies and Jan Sidiangco for his craniotomy surgery expertise.

## Movies

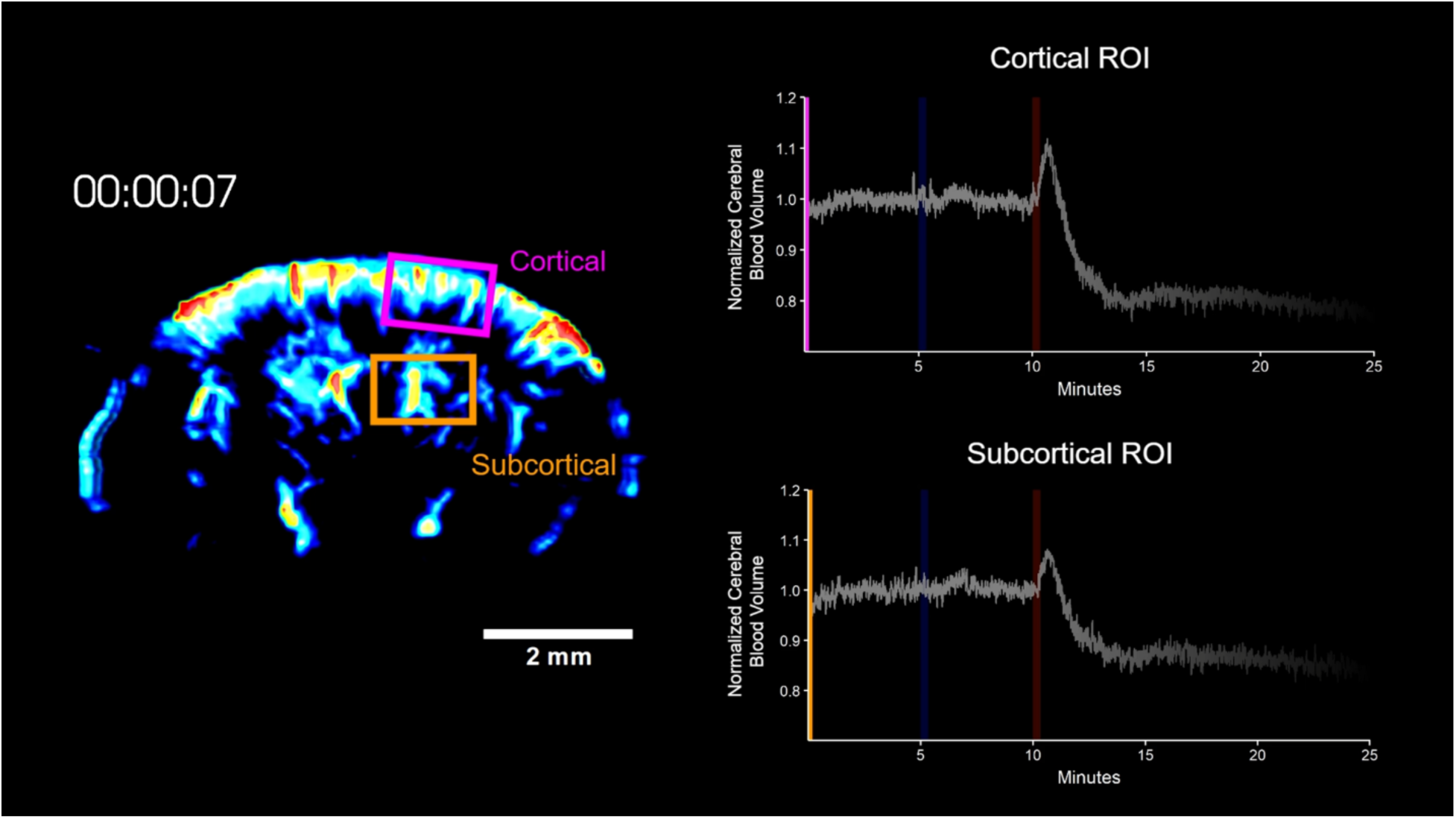

Movie 1. Rose Bengal reduces cerebral blood volume. Example of fUS imaging of an I.P PBS injection (blue vertical shading) followed by an I.P RB injection (red vertical shading). ROIs are drawn cortically and subcortically. PBS results in no appreciable change to blood volume while RB results in a drop across the entire brain.

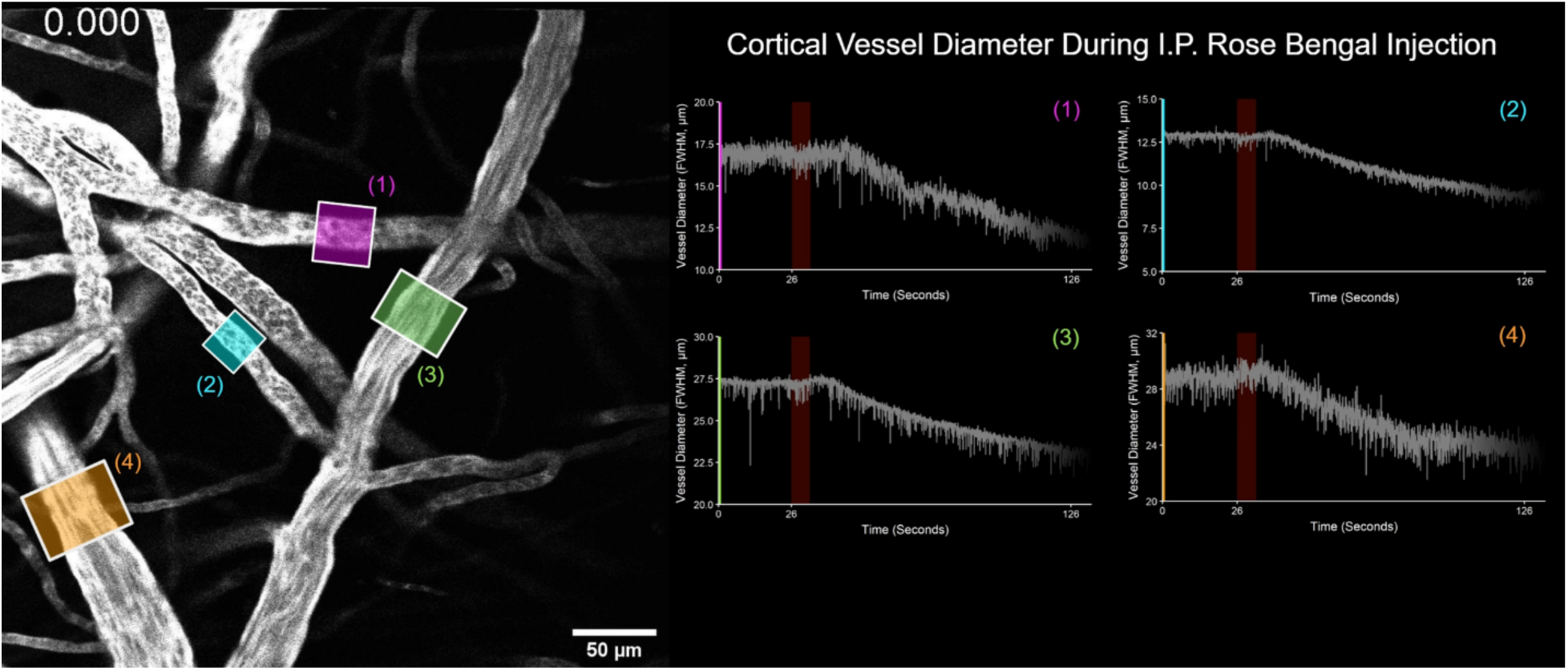

Movie 2. Rose Bengal constricts cortical blood vessels. Example of FITC-Dextran (70kDa) labelled blood vessels before, during, and after the I.P injection of RB. RB is injected starting at 26s over the course of 10 seconds (red shading in the plots). Regions of interest are drawn over vessel segments, and the average diameter (full width at half maximum (FWHM) intensity) of each corresponding vessel is plotted on the right. Vessel diameter decreases to 85% within 100 seconds of injection, a vasoconstriction which corresponds to the cross-sectional area of the vessel reducing to ∼70%. Note the behaviour of vessel #1, where what appears to be some obstruction is flushed through and the flow direction is reversed.

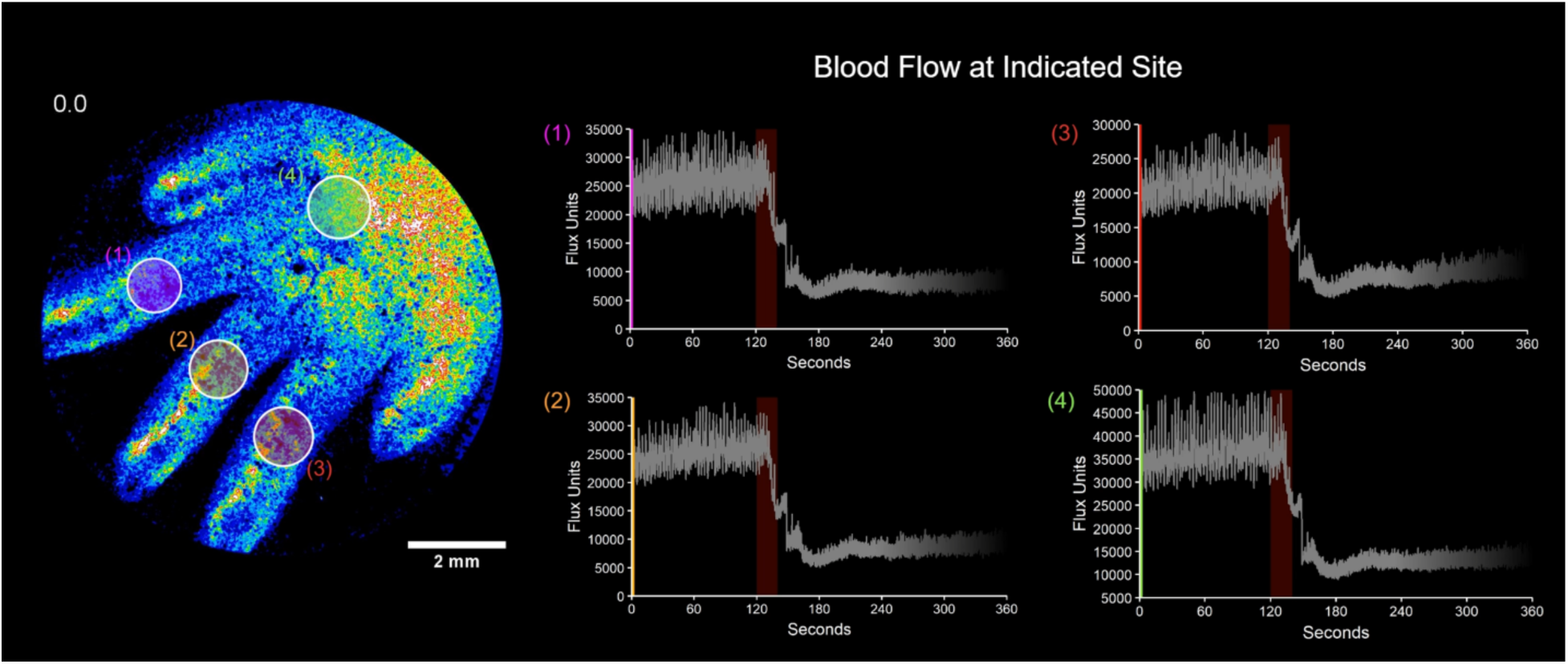

Movie 3. Rose Bengal reduces peripheral blood flow. Example of LSCI of the hind paw during an I.P RB injection (red vertical shading). ROIs show blood flow in different regions of the paw and indicate hypoemia following injection at 120s.

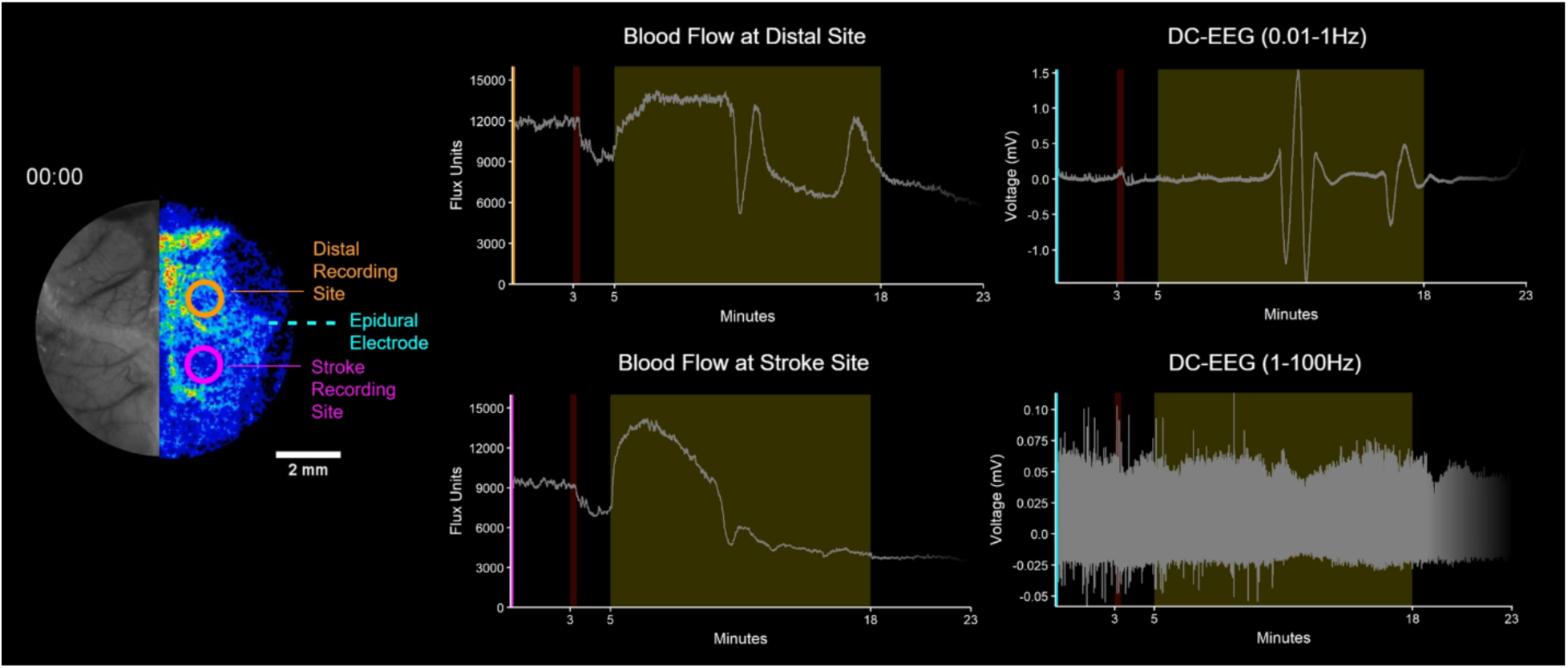

Movie 4. The hemodynamic and electrophysiological monitoring of yellow light (561nm) photothrombosis. LSCI during the stroke is projected onto the right hemisphere. A ROI was drawn over the stroke site (magenta) and one more distally within the same hemisphere (orange) and the average pixel intensity (blood flow) for both ROIs is reported on the right. A permanently implanted epidural electrode (blue) was also monitoring this hemisphere during stroke and that trace has been filtered and presented in both its low (0.01-1Hz) and high (1-100Hz) frequency components. Of note, the RB injection takes place at 3 minutes in this recording.

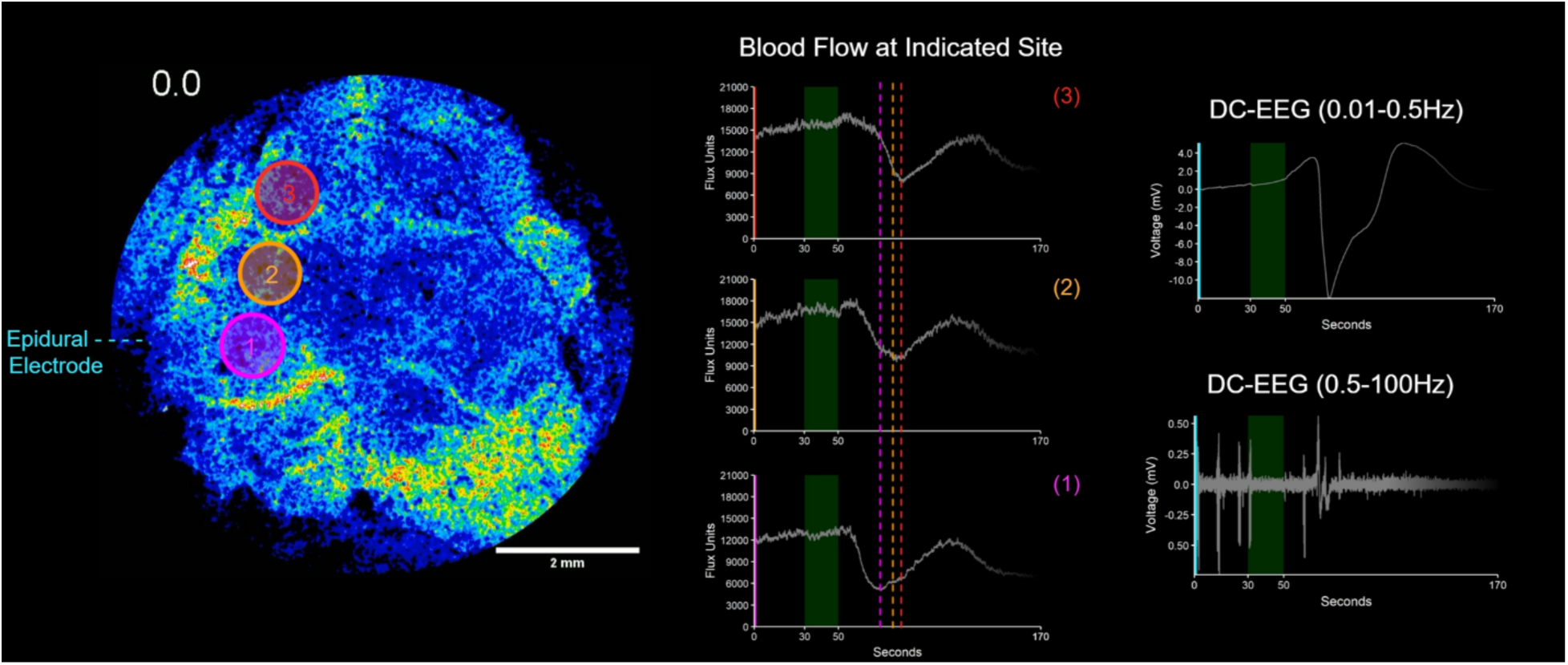

Movie 5. Induction of an optogenetic spreading depolarization with green light (520nm) monitored by LSCI and EEG in a Thy1-ChR2 mouse. Green illumination (lower intensity than during stroke - 3mW/mm^2^) begins at 30 seconds in the left hemisphere (site 1). LSCI ROIs indicate that an ischemic transient propagates outward from site 1 towards the more distal sites (vertical hashed lines indicate the minimum of the ischemic trough for the three sites). The ischemic transient occurs in time with a DC shift and depression of the spontaneous EEG.

## Supplemental Figures

**Figure S1.**
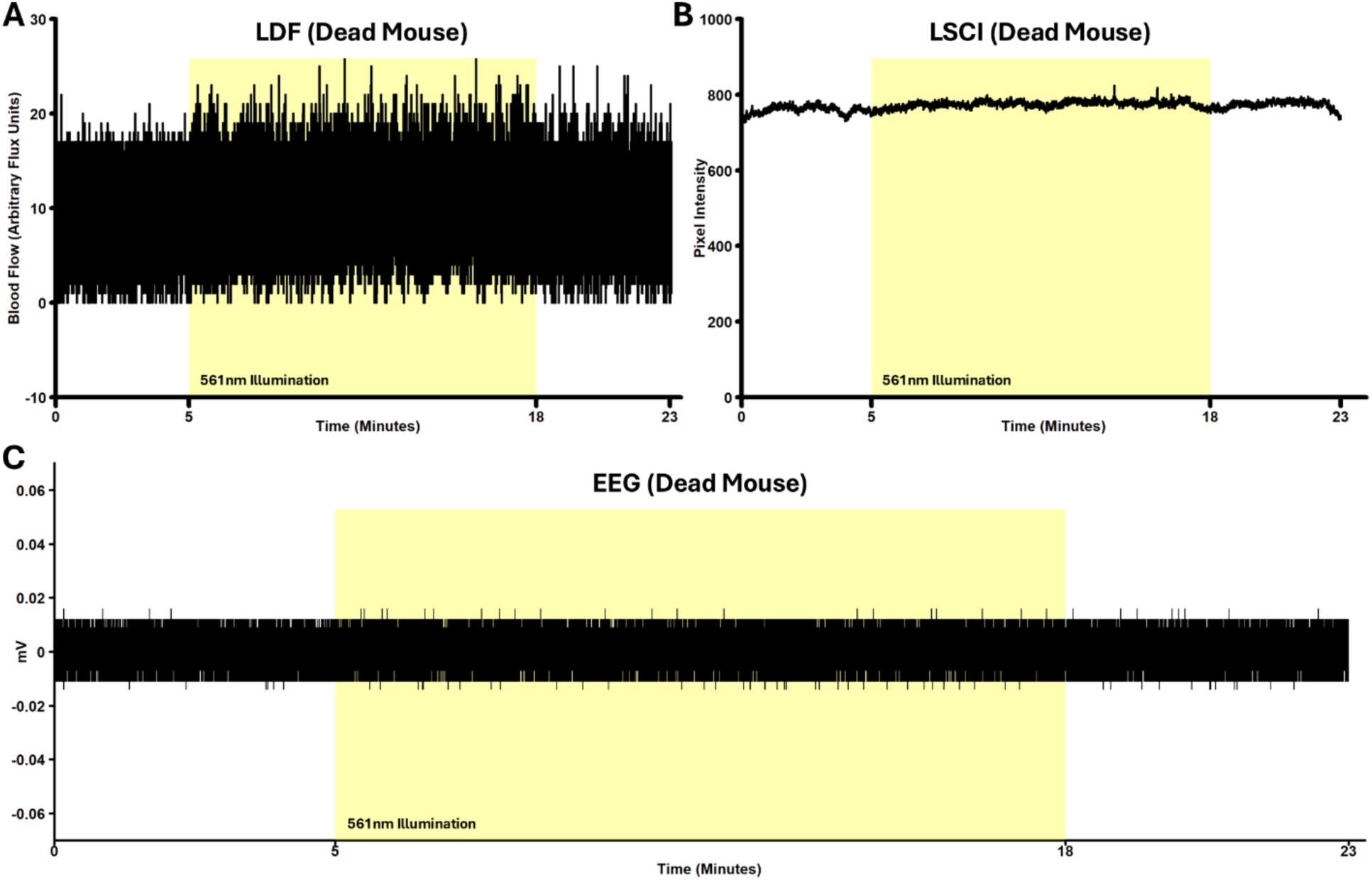
The stimulation light does not crosstalk with LDF, LSCI, or EEG measurements. **A**. Representative example of a transcranial LDF recording of yellow light (561nm) stimulation in a dead mouse (note that LDF blood flow sits at >1000 flux units in a live mouse – Fig.8B). The LDF filters do a sufficient job blocking the 561nm (and 520nm – data not shown) light. **B**. Representative example of a transcranial LSCI recording of yellow light (561nm) stimulation in a dead mouse (ROI drawn over the stimulation site). The LSCI filters do a sufficient job blocking the 561nm (and 520nm – data not shown) light. **C**. Representative example of a DC-EEG (0.01-100Hz) recording of yellow light (561nm) stimulation in a dead mouse. The photothrombotic light has no influence on EEG (no evidence of any photoelectric effects).

**Figure S2.**
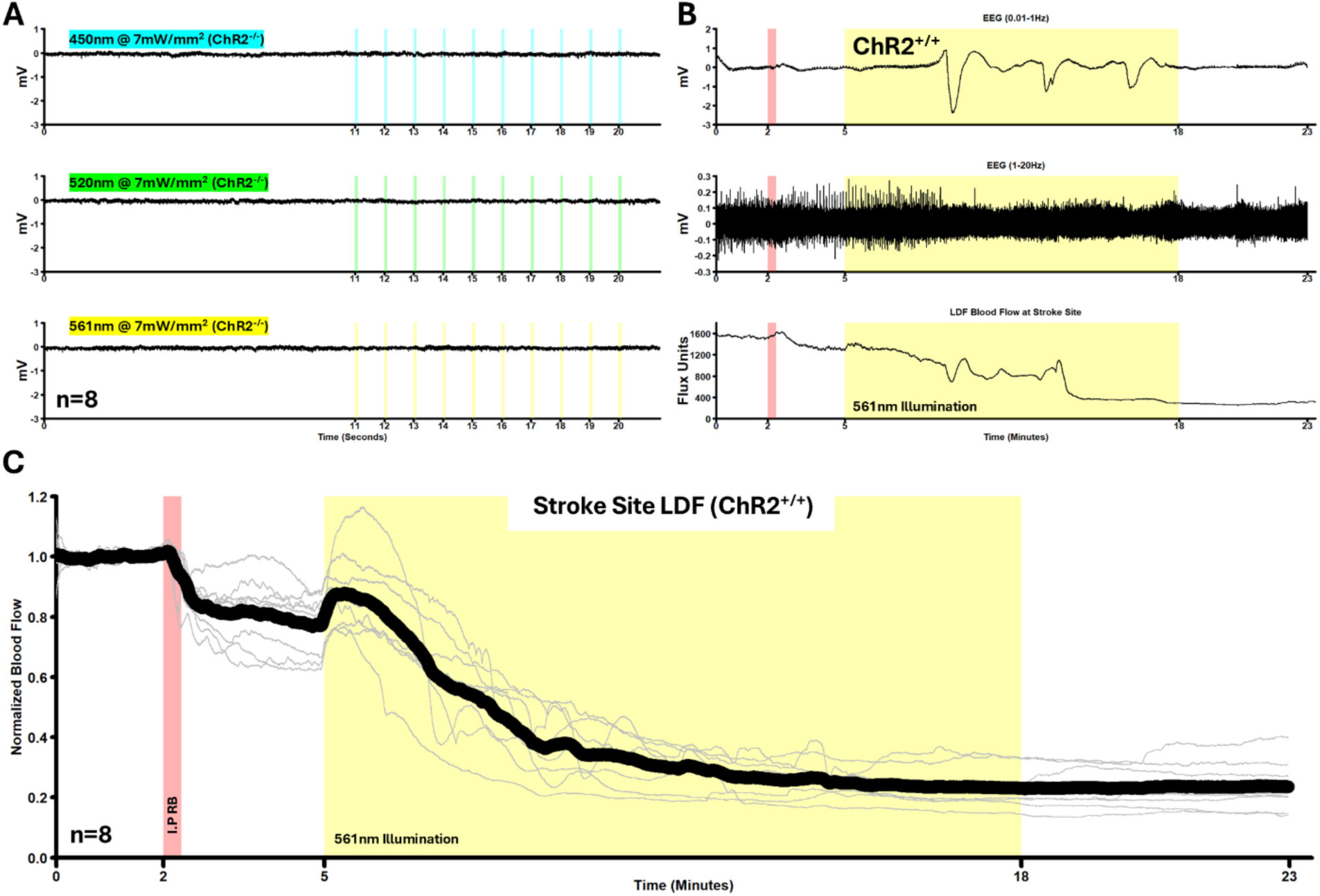
The Crosstalk between photothrombosis and ChR2. **A**. Blue (450nm), green (520nm), and yellow (561nm) light-evoked EEG deflections in ChR2 negative mice. 20ms (interpulse interval of 980ms) light pulses were delivered and the traces from 8 mice were averaged together. **B**. Representative example of aligned EEG and LDF traces from one ChR2 expressing mouse during yellow light (561nm) photothrombosis. EEG is filtered into its low (DC shifts) and high frequency (general cortical activity) components. **C**. Normalized LDF recorded through the transcranial window during yellow light (561nm) photothrombosis in ChR2 expressing mice. The infrared laser was targeted within the centre of the photothrombotic laser, giving a measurement of cortical blood flow in the core of the stroke. The red vertical shading indicates the injection of RB where a hind paw was lifted and an I.P injection was made. The yellow shading indicates the time for which the photothrombosis laser was on (561nm @ 7mW/mm^2^ @ 2.2mm ⌀). Yellow light strokes are identical between ChR2 expressing and non expressing mice.

**Figure S3.**
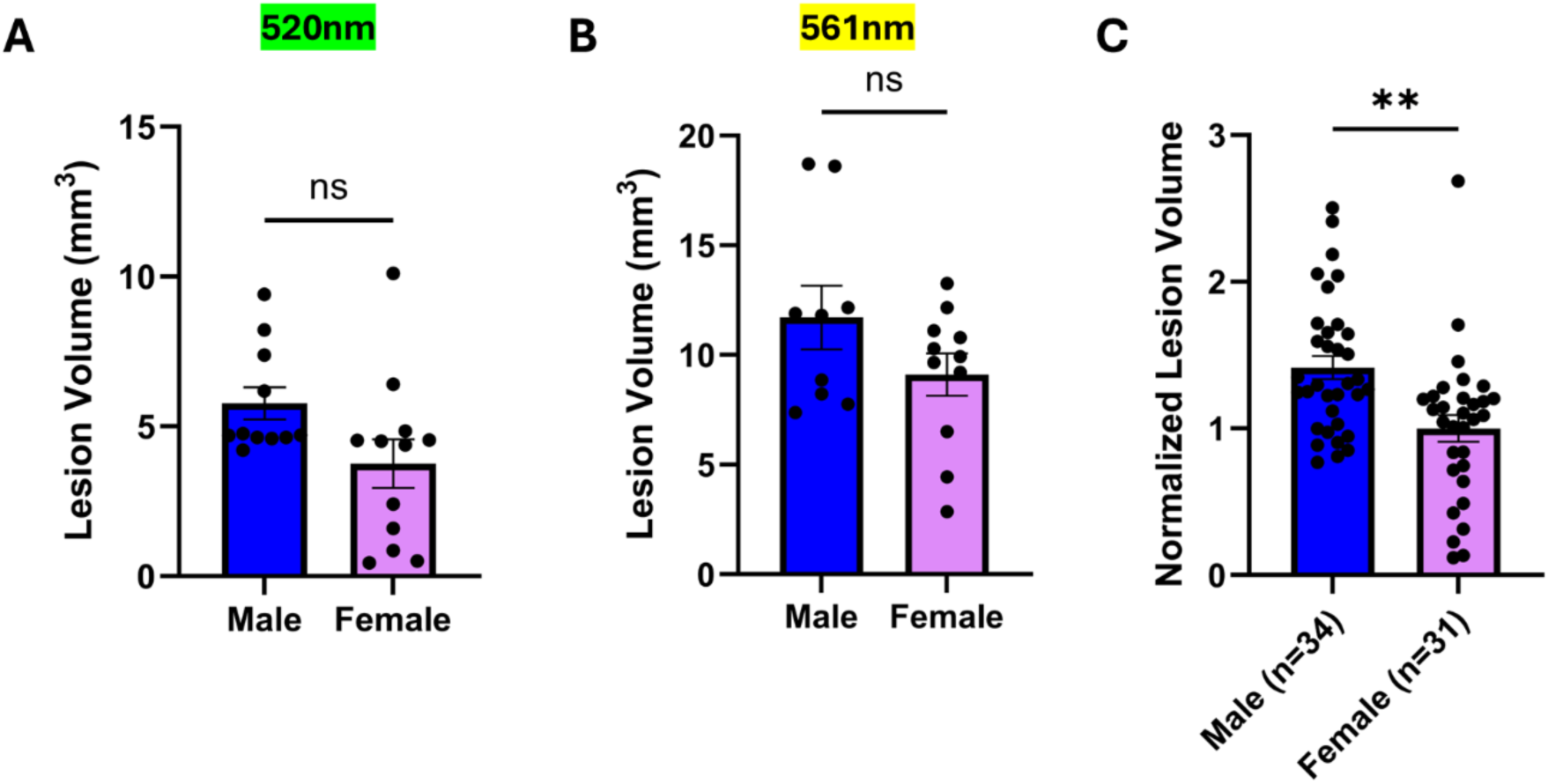
Trend of male strokes being larger than female strokes. **A**. Green (520nm) light lesions collected and measured 7 days post stroke induction (Unpaired Welch’s t = 2.058, df = 18.77, p = 0.0538). **B**. Yellow (561nm) light lesions collected and measured 7 days post stroke induction (Unpaired Welch’s t = 1.490, df = 14.43, p = 0.1578). **C**. Lesion volume from all green (1 cohort) and yellow (2 cohorts) stroke cohorts normalized to average female lesion volume and then pooled together (Unpaired Welch’s t = 3.457, df = 60.79, p = 0.0010).

**Figure S4.**
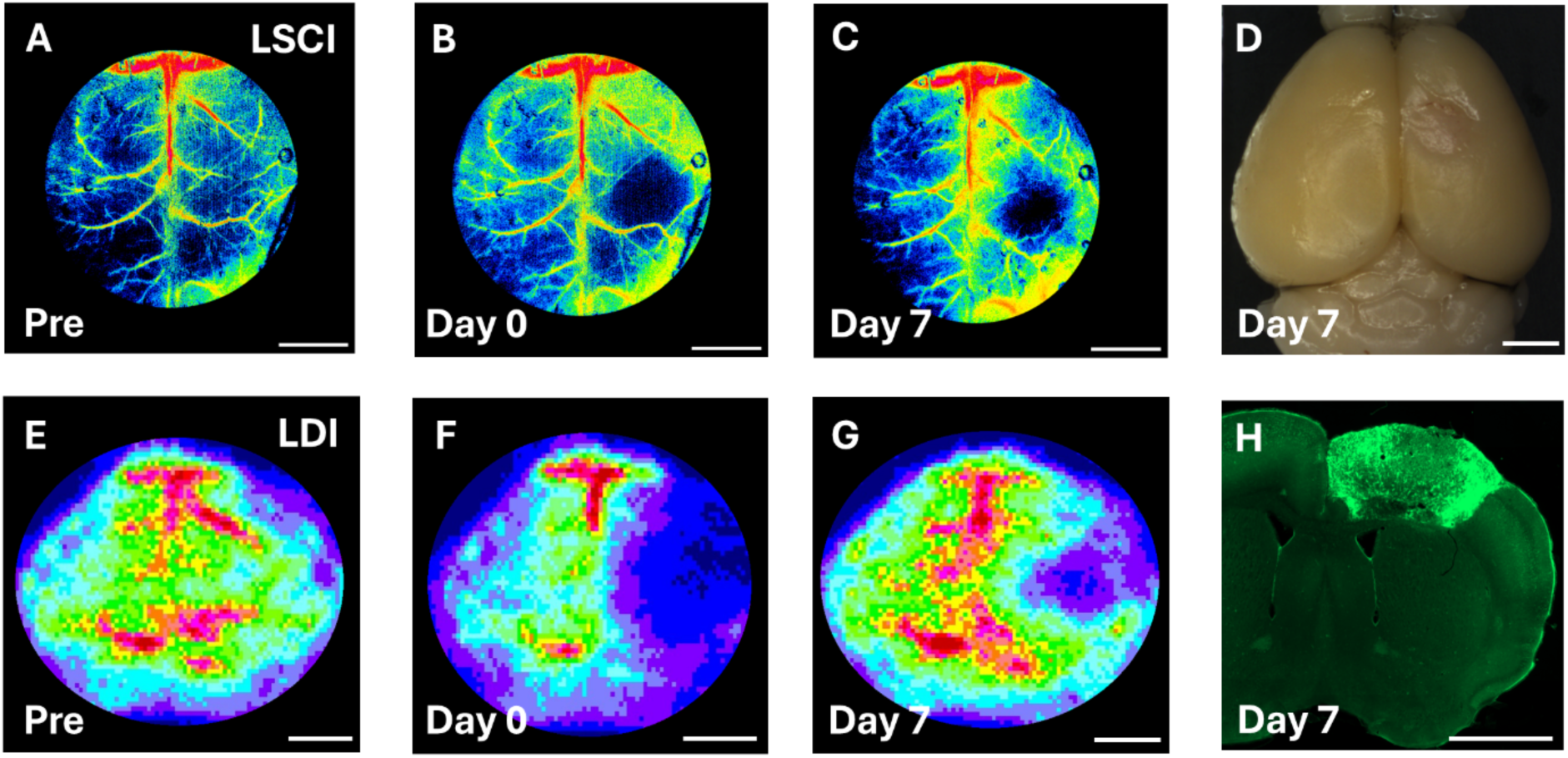
Monitoring of lesion progression by LSCI, LDI, and histology. **A/E**. 5 minutes before induction of the yellow light (561nm) photothrombotic strokes, maps of cortical blood flow were derived by LSCI and LDI through the transcranial window. Such maps were also produced 5 minutes (**B/F**) and 7 days (**C/G**) after induction of the stroke. **D**. Overhead image of the brain showing a photothrombotic stroke at 7 days post injury. **H**. 100µm thick section of brain showing a photothrombotic stroke. Scale bars are 2mm.

**Figure S5.**
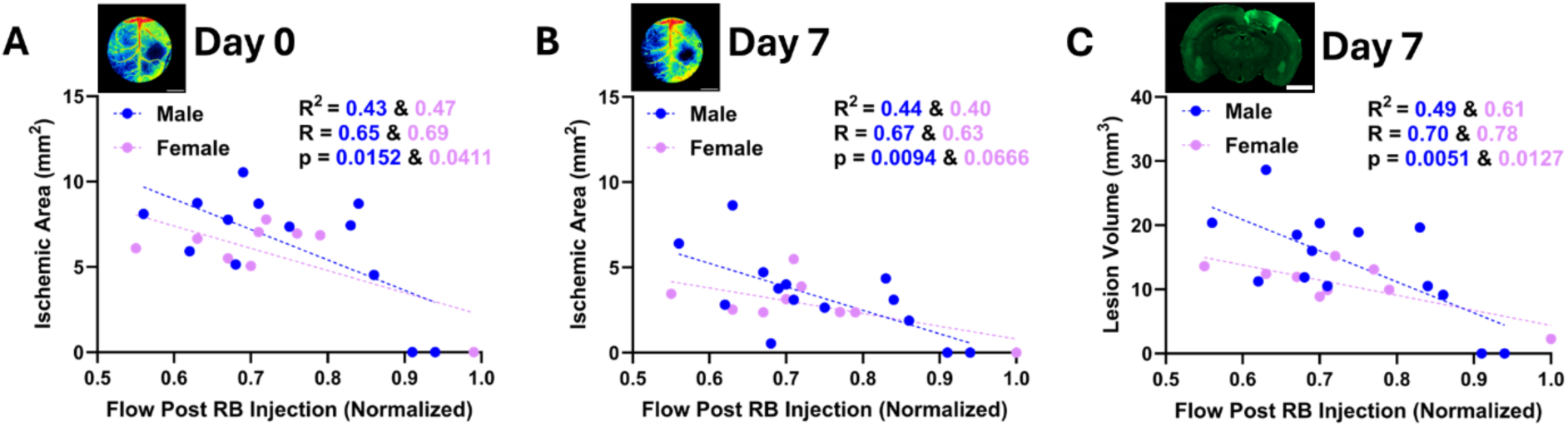
The Rose Bengal induced drop in blood flow partially predicts the size of the photothrombotic lesion. **A**. The LDF measured drop in blood flow just preceding photothrombotic light illumination (4:30-5:00) was corelated against the LSCI determined ischemic area immediately after stroke induction. **B**. The LDF measured drop in blood flow just preceding photothrombotic light illumination (4:30-5:00) was corelated against the LSCI determined ischemic area 7 days after stroke induction. **C**. The LDF measured drop in blood flow just preceding photothrombotic light illumination (4:30-5:00) was corelated against the histology determined lesion volume 7 days after stroke induction. In all cases there appears to be a relationship, however, once the outlier mice who did not have strokes (nor a large RB induced flow drop) are removed, the trends disappear. The interpretation is that a drop in flow following I.P RB indicates a successful injection that will result in a stroke (but does not predict the size of the stroke), whereas a lack of a drop following I.P RB suggests that it was probably a failed injection (bladder, muscle, leg…) and no stroke will be possible.

**Figure S6.**
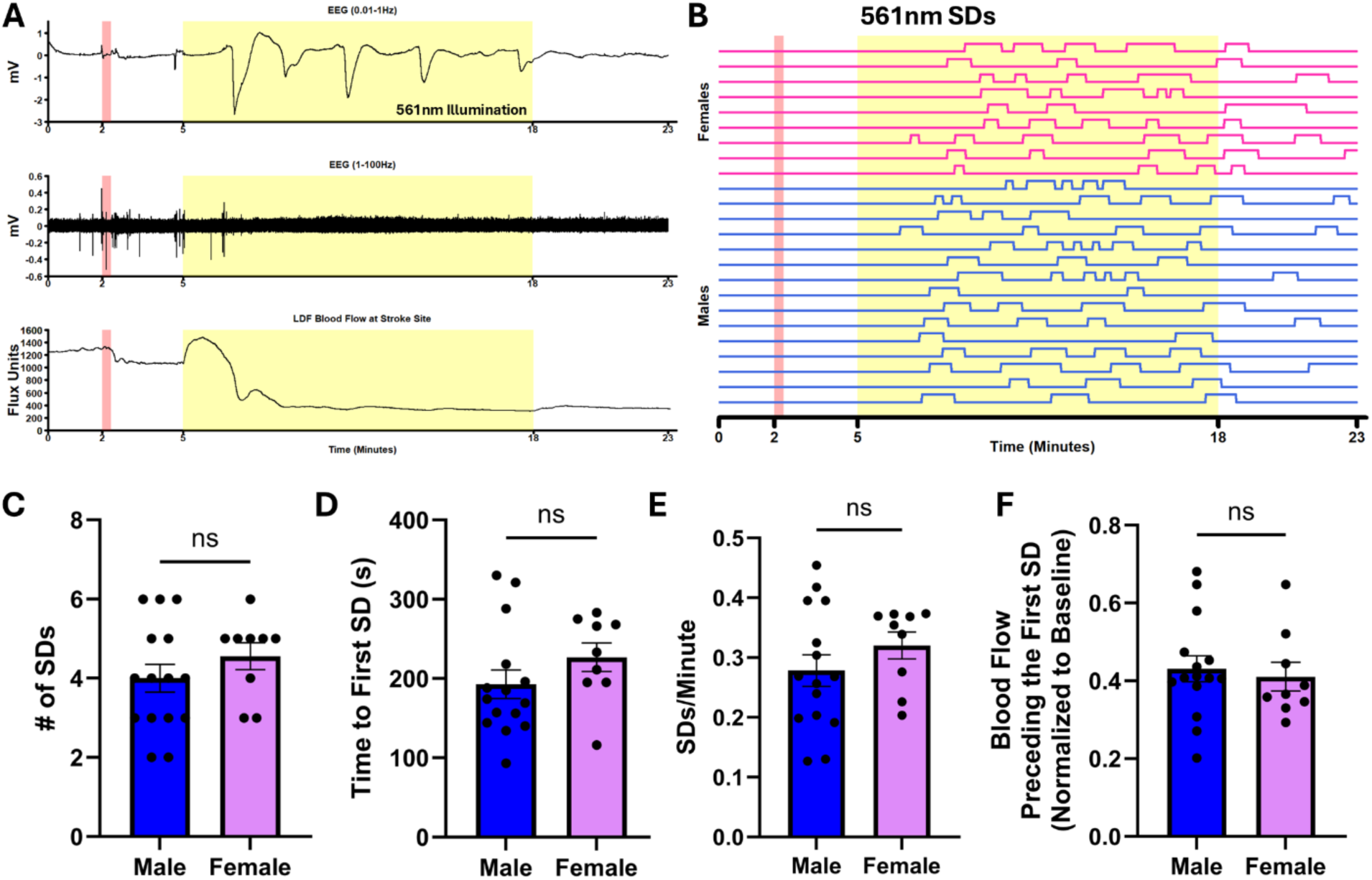
Electrophysiological sex differences in photothrombosis. **A**. Representative example of aligned EEG and LDF traces from one mouse. EEG is filtered into its low (DC shifts) and high frequency (general cortical activity) components. **B**. The occurrence of SDs across all male and female mice. Each SD corresponds to an individual DC shift from the EEG. **C**. The total number of SDs recorded in male and female mice. **D**. The time delay from yellow light illumination to the first SD in male and female mice. **E**. The frequency of SDs calculated from the first one to the end of the recording for each male and female mouse. **F**. The normalized blood flow value (LDF) at which the first SD occurred for both sexes.

## Notes

### Competing Interest Statement

The authors have declared no competing interest.

## References

Abe Y, Kwon S, Oishi M, Unekawa M, Takata N, Seki F, Koyama R, Abe M, Sakimura K, Masamoto K, Tomita Y, Okano H, Mushiake H, Tanaka KF (2021) Optical manipulation of local cerebral blood flow in the deep brain of freely moving mice. Cell Rep 36.

Alarcón E, Edwards AM, Aspée A, Borsarelli CD, Lissi EA (2009) Photophysics and photochemistry of rose bengal bound to human serum albumin. Photochemical and Photobiological Sciences 8:933–943.

Allegra Mascaro AL, Conti E, Lai S, Di Giovanna AP, Spalletti C, Alia C, Panarese A, Scaglione A, Sacconi L, Micera S, Caleo M, Pavone FS (2019) Combined Rehabilitation Promotes the Recovery of Structural and Functional Features of Healthy Neuronal Networks after Stroke. Cell Rep 28:3474–3485.e6.

Allen BD, Singer AC, Boyden ES (2015) Principles of designing interpretable optogenetic behavior experiments. Learning and Memory 22:232–238.

Anenberg E, Arstikaitis P, Niitsu Y, Harrison TC, Boyd JD, Hilton BJ, Tetzlaff W, Murphy TH (2014) Ministrokes in channelrhodopsin-2 transgenic mice reveal widespread deficits in motor output despite maintenance of cortical neuronal excitability. Journal of Neuroscience 34:1094–1104.

Arenkiel BR, Peca J, Davison IG, Feliciano C, Deisseroth K, Augustine GJJ, Ehlers MD, Feng G (2007) In Vivo Light-Induced Activation of Neural Circuitry in Transgenic Mice Expressing Channelrhodopsin-2. Neuron 54:205–218.

Arevalo JF, Garcia RA, Mendoza AJ (2005) Indocyanine green-mediated photothrombosis with intravitreal triamcinolone acetonide for subfoveal choroidal neovascularization in age-related macular degeneration. Graefe’s Archive for Clinical and Experimental Ophthalmology 243:1180–1185.

Arias-Gil G, Ohl FW, Takagaki K, Lippert MT (2016) Measurement, modeling, and prediction of temperature rise due to optogenetic brain stimulation. Neurophotonics 3:045007.

Aswendt M, Pallast N, Wieters F, Baues M, Hoehn M, Fink GR (2021) Lesion Size- and Location-Dependent Recruitment of Contralesional Thalamus and Motor Cortex Facilitates Recovery after Stroke in Mice. Transl Stroke Res 12:87–97.

Balbi M, Vanni MP, Silasi G, Sekino Y, Bolanos L, LeDue JM, Murphy TH (2017) Targeted ischemic stroke induction and mesoscopic imaging assessment of blood flow and ischemic depolarization in awake mice. Neurophotonics 4:1.

Balbi M, Xiao D, Jativa Vega M, Hu H, Vanni MP, Bernier LP, LeDue J, MacVicar B, Murphy TH (2021) Gamma frequency activation of inhibitory neurons in the acute phase after stroke attenuates vascular and behavioral dysfunction. Cell Rep 34.

Baptista MS, Cadet J, Di Mascio P, Ghogare AA, Greer A, Hamblin MR, Lorente C, Nunez SC, Ribeiro MS, Thomas AH, Vignoni M, Yoshimura TM (2017) Type I and Type II Photosensitized Oxidation Reactions: Guidelines and Mechanistic Pathways. Photochem Photobiol 93:912–919.

Bardutzky J, Shen Q, Henninger N, Bouley J, Duong TQ, Fisher M (2005) Differences in ischemic lesion evolution in different rat strains using diffusion and perfusion imaging. Stroke 36:2000–2005.

Behl C, Skutella T, Lezoualc’h F, Post A, Widmann M, Newton CJ, Holsboer F (1997) Neuroprotection against oxidative stress by estrogens: structure-activity relationship. Mol Pharmacol 51:535–541 Available at: http://www.ncbi.nlm.nih.gov/pubmed/9106616.

Beringhs AO, Singh SP, Valdez TA, Lu X (2020) Sublingual indocyanine green films for non-invasive swallowing assessment and inflammation detection through NIR/SWIR optical imaging. Sci Rep 10.

Bice AR, Xiao Q, Kong J, Yan P, Rosenthal ZP, Kraft AW, Smith K, Wieloch T, Lee JM, Culver JP, Bauer AQ (2022) Homotopic contralesional excitation suppresses spontaneous circuit repair and global network reconnections following ischemic stroke. Elife 11.

Bo B, Li Y, Li W, Wang Y, Tong S (2019) Optogenetic Excitation of Ipsilesional Sensorimotor Neurons is Protective in Acute Ischemic Stroke: A Laser Speckle Imaging Study. IEEE Trans Biomed Eng 66:1372–1379.

Bo B, Li Y, Li W, Wang Y, Tong S (2020) Optogenetic translocation of protons out of penumbral neurons is protective in a rodent model of focal cerebral ischemia. Brain Stimul 13:881–890.

Boquillon M, Boquillon JP, Bralet J (1992) Photochemically induced, graded cerebral infarction in the mouse by laser irradiation evolution of brain edema. J Pharmacol Toxicol Methods 27:1–6.

Boyce AKJ, Fouad Y, Gom RC, Ashby DM, Martins-Silva C, Molina L, Füzesi T, Ens C, Nicola W, McGirr A, Teskey GC, Thompson RJ (2025) Contralesional hippocampal spreading depolarization promotes functional recovery after stroke. Nature Communications 16.

Boyden ES, Zhang F, Bamberg E, Nagel G, Deisseroth K (2005) Millisecond-timescale, genetically targeted optical control of neural activity. Nat Neurosci 8:1263–1268.

Brunner C, Grillet M, Urban A, Roska B, Montaldo G, Macé E (2021) Whole-brain functional ultrasound imaging in awake head-fixed mice. Nat Protoc 16:3547–3571 Available at: 10.1038/s41596-021-00548-8.

Busto R, Dietrich WD, Globus MY, Ginsberg MD (1989) The importance of brain temperature in cerebral ischemic injury. Stroke 20:1113–1114 Available at: https://www.ahajournals.org/doi/10.1161/01.STR.20.8.1113.

Busto R, Dietrich WD, Globus MY-T, Valdés I, Scheinberg P, Ginsberg MD (1987) Small Differences in Intraischemic Brain Temperature Critically Determine the Extent of Ischemic Neuronal Injury. Journal of Cerebral Blood Flow & Metabolism 7:729–738 Available at: https://journals.sagepub.com/doi/10.1038/jcbfm.1987.127.

Cai H, Yao H, Ibayashi S, Uchimura H, Fujishima M (1998) Photothrombotic Middle Cerebral Artery Occlusion in Spontaneously Hypertensive Rats: Influence of Substrain, Gender, and Distal Middle Cerebral Artery Patterns on Infarct Size. Stroke 29:1982–1987 Available at: https://www.ahajournals.org/doi/10.1161/01.STR.29.9.1982.

Canals S, Makarova I, López-Aguado L, Largo C, Ibarz JM, Herreras O (2005) Longitudinal depolarization gradients along the somatodendritic axis of CA1 pyramidal cells: A novel feature of spreading depression. J Neurophysiol 94:943–951.

Carmichael ST (2005) Rodent Models of Focal Stroke: Size, Mechanism, and Purpose. NeuroRX 2:396–409.

Christie IN, Wells JA, Southern P, Marina N, Kasparov S, Gourine A V., Lythgoe MF (2013) FMRI response to blue light delivery in the naïve brain: Implications for combined optogenetic fMRI studies. Neuroimage 66:634–641.

Clark TA, Sullender C, Kazmi SM, Speetles BL, Williamson MR, Palmberg DM, Dunn AK, Jones TA (2019) Artery targeted photothrombosis widens the vascular penumbra, instigates peri-infarct neovascularization and models forelimb impairments. Sci Rep 9.

Conti E, Scaglione A, de Vito G, Calugi F, Pasquini M, Pizzorusso T, Micera S, Allegra Mascaro AL, Pavone FS (2022) Combining Optogenetic Stimulation and Motor Training Improves Functional Recovery and Perilesional Cortical Activity. Neurorehabil Neural Repair 36:107–118.

Corbett D, Carmichael ST, Murphy TH, Jones TA, Schwab ME, Jolkkonen J, Clarkson AN, Dancause N, Weiloch T, Johansen-Berg H, Nilsson M, McCullough LD, Joy MT (2017) Enhancing the Alignment of the Preclinical and Clinical Stroke Recovery Research Pipeline: Consensus-Based Core Recommendations from the Stroke Recovery and Rehabilitation Roundtable Translational Working Group ∗. Neurorehabil Neural Repair 31:699–707.

Costa RA, Scapucin L, Moraes NS, Calucci D, Melo LA, Cardillo JA, Farah ME (2002) Indocyanine green-mediated photothrombosis as a new technique of treatment for persistent central serous chorioretinopathy. Curr Eye Res 25:287–297.

Cotrina ML, Lou N, Tome-Garcia J, Goldman J, Nedergaard M (2017) Direct comparison of microglial dynamics and inflammatory profile in photothrombotic and arterial occlusion evoked stroke. Neuroscience 343:483–494 Available at: 10.1016/j.neuroscience.2016.12.012.

Delorme A, Makeig S (2004) EEGLAB: An open source toolbox for analysis of single-trial EEG dynamics including independent component analysis. J Neurosci Methods 134:9– 21.

Devor A, Hillman EMC, Tian P, Waeber C, Teng IC, Ruvinskaya L, Shalinsky MH, Zhu H, Haslinger RH, Narayanan SN, Ulbert I, Dunn AK, Lo EH, Rosen BR, Dale AM, Kleinfeld D, Boas DA (2008) Stimulus-induced changes in blood flow and 2-deoxyglucose uptake dissociate in ipsilateral somatosensory cortex. Journal of Neuroscience 28:14347–14357.

Dreier JP et al. (2017) Recording, analysis, and interpretation of spreading depolarizations in neurointensive care: Review and recommendations of the COSBID research group. Journal of Cerebral Blood Flow and Metabolism 37:1595–1625.

Dreier JP, Major S, Lemale CL, Kola V, Reiffurth C, Schoknecht K, Hecht N, Hartings JA, Woitzik J (2019) Correlates of spreading depolarization, spreading depression, and negative ultraslow potential in epidural versus subdural electrocorticography. Front Neurosci 13.

Dula AN, Mlynash M, Zuck ND, Albers GW, Warach SJ (2020) Neuroimaging in Ischemic Stroke Is Different Between Men and Women in the DEFUSE 3 Cohort. Stroke 51:481– 488.

Errico C, Pierre J, Pezet S, Desailly Y, Lenkei Z, Couture O, Tanter M (2015) Ultrafast ultrasound localization microscopy for deep super-resolution vascular imaging. Nature 527:499–502.

Farah ME, Cardillo JA, Luzardo AC, Calucci D, Williams GA, Costa RA (2004) Indocyanine green mediated photothrombosis for the management of predominantly classic choroidal neovascularisation caused by age related macular degeneration. British Journal of Ophthalmology 88:1055–1059.

Fenno LE, Gunaydin LA, Deisseroth K (2015) Mapping anatomy to behavior in Thy1:18 ChR2-YFP transgenic mice using optogenetics. Cold Spring Harb Protoc 2015:537– 548.

Fluri F, Schuhmann MK, Kleinschnitz C (2015) Animal models of ischemic stroke and their application in clinical research. Drug Des Devel Ther 9:3445–3454.

Futrell N (1991) An improved photochemical model of embolic cerebral infarction in rats. Stroke 22:225–232 Available at: https://www.ahajournals.org/doi/abs/10.1161/01.STR.22.2.225.

Ginsberg MD, Busto R (1989) Rodent models of cerebral ischemia. Stroke 20:1627–1642 Available at: https://www.ahajournals.org/doi/10.1161/01.STR.20.12.1627.

Harrison TC, Silasi G, Boyd JD, Murphy TH (2013) Displacement of Sensory Maps and Disorganization of Motor Cortex After Targeted Stroke in Mice. Stroke 44:2300–2306 Available at: 10.1161/STROKEAHA.113.001272.

Hartmann DA, Berthiaume AA, Grant RI, Harrill SA, Koski T, Tieu T, McDowell KP, Faino A V., Kelly AL, Shih AY (2021) Brain capillary pericytes exert a substantial but slow influence on blood flow. Nat Neurosci 24:633–645.

Henninger N, Sicard KM, Schmidt KF, Bardutzky J, Fisher M (2006) Comparison of ischemic lesion evolution in embolic versus mechanical middle cerebral artery occlusion in Sprague Dawley rats using diffusion and perfusion imaging. Stroke 37:1283–1287.

Herrmann KS (1983) Platelet aggregation induced in the hamster cheek pouch by a photochemical process with excited fluorescein isothiocyanate-dextran. Microvasc Res 26:238–249 Available at: https://linkinghub.elsevier.com/retrieve/pii/0026286283900730.

Hilger T, Blunk JA, Hoehn M, Mies G, Wester P (2004) Characterization of a Novel Chronic Photothrombotic Ring Stroke Model in Rats by Magnetic Resonance Imaging, Biochemical Imaging, and Histology. Journal of Cerebral Blood Flow & Metabolism 24:789–797 Available at: 10.1097/01.WCB.0000123905.17746.DB.

Hill RA, Tong L, Yuan P, Murikinati S, Gupta S, Grutzendler J (2015) Regional Blood Flow in the Normal and Ischemic Brain Is Controlled by Arteriolar Smooth Muscle Cell Contractility and Not by Capillary Pericytes. Neuron 87:95–110.

Hille B (2001) Ion channels of excitable membranes, 3rd ed. Sunderland, Mass: Sinauer.

Hodgkin AL, Katz B (1949) The effect of temperature on the electrical activity of the giant axon of the squid. J Physiol 109:240–249 Available at: http://www.ncbi.nlm.nih.gov/pubmed/15394322.

Ishikawa M, Sekizuka E, Oshio C, Sato S, Yamaguchi N, Terao S, Tsukada K, Minamitani H, Kawase T (2002) Platelet adhesion and arteriolar dilation in the photothrombosis: observation with the rat closed cranial and spinal windows. J Neurol Sci 194:59–69 Available at: https://www.sciencedirect.com/science/article/pii/S0022510X01006736.

Kaiser T, Feng G (2019) Tmem119-EGFP and Tmem119-creERT2 transgenic mice for labeling and manipulating microglia. eNeuro 6:1–18.

Kam CY, Singh ID, Gonzalez DG, Matte-Martone C, Solá P, Solanas G, Bonjoch J, Marsh E, Hirschi KK, Greco V (2023) Mechanisms of skin vascular maturation and maintenance captured by longitudinal imaging of live mice. Cell 186:2345–2360.e16.

Kim Y, Busto R, Dietrich WD, Kraydieh S, Ginsberg MD (1996) Delayed postischemic hyperthermia in awake rats worsens the histopathological outcome of transient focal cerebral ischemia. Stroke 27:2274–2280; discussion 2281 Available at: http://www.ncbi.nlm.nih.gov/pubmed/8969793.

Knezic A, Broughton BRS, Widdop RE, McCarthy CA (2022) Optimising the photothrombotic model of stroke in the C57BI/6 and FVB/N strains of mouse. Sci Rep 12.

Kuroiwa T, Xi G, Hua Y, Nagaraja TN, Fenstermacher JD, Keep RF (2009) Development of a rat model of photothrombotic ischemia and infarction within the caudoputamen. Stroke 40:248–253.

Labat-gest V, Tomasi S (2013) Photothrombotic ischemia: a minimally invasive and reproducible photochemical cortical lesion model for mouse stroke studies. J Vis Exp 76.

Lee J-K, Park M-S, Kim Y-S, Moon K-S, Joo S-P, Kim T-S, Kim J-H, Kim S-H (2007) Photochemically induced cerebral ischemia in a mouse model. Surg Neurol 67:620– 625; discussion 625 Available at: http://www.ncbi.nlm.nih.gov/pubmed/17512331.

Leng C, Li Y, Sun Y, Liu W (2023) Induction of Acute Ischemic Stroke in Mice Using the Distal Middle Artery Occlusion Technique. Journal of Visualized Experiments 2023.

Li DY, Xia Q, Yu TT, Zhu JT, Zhu D (2021) Transmissive-detected laser speckle contrast imaging for blood flow monitoring in thick tissue: from Monte Carlo simulation to experimental demonstration. Light Sci Appl 10.

Li H, Zhang N, Lin H-Y, Yu Y, Cai Q-Y, Ma L, Ding S (2014) Histological, cellular and behavioral assessments of stroke outcomes after photothrombosis-induced ischemia in adult mice. BMC Neurosci 15:58 Available at: http://www.ncbi.nlm.nih.gov/pubmed/24886391.

Lim DH, Ledue JM, Mohajerani MH, Murphy TH (2014) Optogenetic mapping after stroke reveals network-wide scaling of functional connections and heterogeneous recovery of the peri-infarct. Journal of Neuroscience 34:16455–16466.

Lin JY (2011) A user’s guide to channelrhodopsin variants: Features, limitations and future developments. Exp Physiol 96:19–25.

Lin Y, Yao M, Wu H, Wu F, Cao S, Ni H, Dong J, Yang D, Sun Y, Kou X, Li J, Xiao H, Chang L, Wu J, Liu Y, Luo C, Zhu D (2021) Environmental enrichment implies GAT-1 as a potential therapeutic target for stroke recovery. Theranostics 11:3760–3780.

Llovera G, Roth S, Plesnila N, Veltkamp R, Liesz A (2014) Modeling stroke in mice: Permanent coagulation of the distal middle cerebral artery. Journal of Visualized Experiments.

Lyu J, Liu L, Guo M, Li S, Su W, Liu J, Ji X (2025) Brain-targeted mild hypothermia ameliorates ischemic brain injury and promotes stroke recovery in aged mice. Journal of Cerebral Blood Flow and Metabolism.

Macé E, Montaldo G, Cohen I, Baulac M, Fink M, Tanter M (2011) Functional ultrasound imaging of the brain. Nat Methods 8:662–664.

Marks K, Ahn SJ, Rai N, Anfray A, Iadecola C, Anrather J (2025) A minimally invasive thrombotic model to study stroke in awake mice. Nature Communications 16.

Matsuno H, Uematsu T, Umemura K, Takiguchi Y, Asai Y, Muranaka Y, Nakashima M (1993) A simple and reproducible cerebral thrombosis model in rats induced by a photochemical reaction and the effect of a plasminogen-plasminogen activator chimera in this model. J Pharmacol Toxicol Methods 29:165–173 Available at: https://linkinghub.elsevier.com/retrieve/pii/105687199390068P.

Mattis J, Tye KM, Ferenczi EA, Ramakrishnan C, O’Shea DJ, Prakash R, Gunaydin LA, Hyun M, Fenno LE, Gradinaru V, Yizhar O, Deisseroth K (2012) Principles for applying optogenetic tools derived from direct comparative analysis of microbial opsins. Nat Methods 9:159–172.

McDonald MW, Dykes A, Jeffers MS, Carter A, Nevins R, Ripley A, Silasi G, Corbett D (2021) Remote Ischemic Conditioning and Stroke Recovery. Neurorehabil Neural Repair 35:545–549.

McDowell KP, Berthiaume AA, Tieu T, Hartmann DA, Shih AY (2021) VasoMetrics: Unbiased spatiotemporal analysis of microvascular diameter in multi-photon imaging applications. Quant Imaging Med Surg 11:969–982.

Mergenthaler P, Meisel A (2012) Do stroke models model stroke? DMM Disease Models and Mechanisms 5:718–725.

Mirza Agha B, Akbary R, Ghasroddashti A, Nazari-Ahangarkolaee M, Whishaw IQ, Mohajerani MH (2021) Cholinergic upregulation by optogenetic stimulation of nucleus basalis after photothrombotic stroke in forelimb somatosensory cortex improves endpoint and motor but not sensory control of skilled reaching in mice. Journal of Cerebral Blood Flow and Metabolism 41:1608–1622.

Navajas E V., Costa RA, Farah ME, Cardillo JA, Bonomo PP (2003) Indocyanine green-mediated photothrombosis combined with intravitreal triamcinolone for the treatment of choroidal neovascularization in serpiginous choroiditis. Eye 17:563–566.

Neckers DC (1989) Rose Bengal. J Photochem Photobiol A Chem 47:1–29 Available at: https://linkinghub.elsevier.com/retrieve/pii/1010603089850026.

Newton JM, Pushie MJ, Sylvain NJ, Hou H, Weese Maley S, Kelly ME (2022) Sex differences in the mouse photothrombotic stroke model investigated with X-ray fluorescence microscopy and Fourier transform infrared spectroscopic imaging. IBRO Neurosci Rep 13:127–135.

Okabe N, Wei X, Abumeri F, Batac J, Hovanesyan M, Dai W, Azarapetian S, Campagna J, Pilati N, Marasco A, Alvaro G, Gunthorpe MJ, Varghese J, Cramer SC, Mody I, Carmichael ST (2025) Parvalbumin interneurons regulate rehabilitation-induced functional recovery after stroke and identify a rehabilitation drug. Nat Commun 16:2556 Available at: https://www.nature.com/articles/s41467-025-57860-0.

Owais SB, Bulwa ZB, Ammar F El (2024) Differences in stroke clinical presentation among sexes. Journal of Stroke and Cerebrovascular Diseases 33.

Owen SF, Liu MH, Kreitzer AC (2019) Thermal constraints on in vivo optogenetic manipulations. Nat Neurosci 22:1061–1065 Available at: 10.1038/s41593-019-0422-3.

Pallikaris IG, Tslimbaris MK, Iliaki OE, Naoumidi II, Georgiades A, Panagopoulos IA (1993) Effectiveness of corneal neovascularization photothrombosis using phthalocyanine and a diode laser (675 nm). Lasers Surg Med 13:197–203.

Pan ZH, Ganjawala TH, Lu Q, Ivanova E, Zhang Z (2014) ChR2 mutants at L132 and T159 with improved operational light sensitivity for vision restoration. PLoS One 9.

Paz JT, Davidson TJ, Frechette ES, Delord B, Parada I, Peng K, Deisseroth K, Huguenard JR (2013) Closed-loop optogenetic control of thalamus as a tool for interrupting seizures after cortical injury. Nat Neurosci 16:64–70.

Peixoto HM, Cruz RMS, Moulin TC, Leão RN (2020) Modeling the Effect of Temperature on Membrane Response of Light Stimulation in Optogenetically-Targeted Neurons. Front Comput Neurosci 14:1–14.

Podgorski K, Ranganathan G (2016) Brain heating induced by near-infrared lasers during multiphoton microscopy. J Neurophysiol 116:1012–1023.

Raman-Nair J, Cron G, MacLeod K, Lacoste B (2023) Sex specific acute cerebrovascular responses to photothrombotic stroke in mice. eNeuro 11:ENEURO.0400-22.2023.

Reglodi D, Somogyvari-Vigh A, Maderdrut JL, Vigh S, Arimura A (2000) Postischemic spontaneous hyperthermia and its effects in middle cerebral artery occlusion in the rat. Exp Neurol 163:399–407.

Rexrode KM, Madsen TE, Yu AYX, Carcel C, Lichtman JH, Miller EC (2022) The Impact of Sex and Gender on Stroke. Circ Res 130:512–528.

Risher WC, Ard D, Yuan J, Kirov SA (2010) Recurrent spontaneous spreading depolarizations facilitate acute dendritic injury in the ischemic penumbra. Journal of Neuroscience 30:9859–9868.

Rosenblum WI, El-Sabban F (1977) Platelet aggregation in the cerebral microcirculation: effect of aspirin and other agents. Circ Res 40:320–328.

Rungta RL, Osmanski BF, Boido D, Tanter M, Charpak S (2017) Light controls cerebral blood flow in naive animals. Nat Commun 8.

Schaffer CB, Friedman B, Nishimura N, Schroeder LF, Tsai PS, Ebner FF, Lyden PD, Kleinfeld D (2006) Two-photon imaging of cortical surface microvessels reveals a robust redistribution in blood flow after vascular occlusion. PLoS Biol 4:258–270.

Schmidt A, Hoppen M, Strecker J-K, Diederich K, Schäbitz W-R, Schilling M, Minnerup J (2012) Photochemically induced ischemic stroke in rats. Exp Transl Stroke Med 4:13.

Schoknecht K, Kikhia M, Lemale CL, Liotta A, Lublinsky S, Mueller S, Boehm-Sturm P, Friedman A, Dreier JP (2021) The role of spreading depolarizations and electrographic seizures in early injury progression of the rat photothrombosis stroke model. Journal of Cerebral Blood Flow and Metabolism 41:413–430.

Shah AM, Ishizaka S, Cheng MY, Wang EH, Bautista AR, Levy S, Smerin D, Sun G, Steinberg GK (2017) Optogenetic neuronal stimulation of the lateral cerebellar nucleus promotes persistent functional recovery after stroke. Sci Rep 7:1–11.

Silasi G, Boyd JD, LeDue J, Murphy TH (2013) Improved methods for chronic light-based motor mapping in mice: Automated movement tracking with accelerometers, and chronic EEG recording in a bilateral thin-skull preparation. Front Neural Circuits 7.

Silasi G, Xiao D, Vanni MP, Chen ACN, Murphy TH (2016) Intact skull chronic windows for mesoscopic wide-field imaging in awake mice. J Neurosci Methods 267:141–149.

Simpkins JW, Singh M, Brock C, Etgen AM (2012) Neuroprotection and estrogen receptors. Neuroendocrinology 96:119–130.

Sinigaglia-Coimbra R, Cavalheiro EA, Coimbra CG (2002) Postischemic hyperthermia induces Alzheimer-like pathology in the rat brain. Acta Neuropathol 103:444–452.

Stujenske JM, Spellman T, Gordon JA (2015) Modeling the Spatiotemporal Dynamics of Light and Heat Propagation for In Vivo Optogenetics. Cell Rep 12:525–534.

Suo Q, Deng L, Chen T, Wu S, Qi L, Liu Z, He T, Tian H-L, Li W, Tang Y, Yang G-Y, Zhang Z (2023) Optogenetic Activation of Astrocytes Reduces Blood-Brain Barrier Disruption via IL-10 In Stroke. Aging Dis 14:1870 Available at: https://www.aginganddisease.org/EN/10.14336/AD.2023.0226.

Tang GL, Kim KJ (2021) Laser doppler perfusion imaging in the mouse hindlimb. Journal of Visualized Experiments 2021.

Tang T, Hu L, Liu Y, Fu X, Li J, Yan F, Cao S, Chen G (2022) Sex-Associated Differences in Neurovascular Dysfunction During Ischemic Stroke. Front Mol Neurosci 15 Available at: https://www.frontiersin.org/articles/10.3389/fnmol.2022.860959/full.

Tennant KA, Taylor SL, White ER, Brown CE (2017) Optogenetic rewiring of thalamocortical circuits to restore function in the stroke injured brain. Nat Commun 8.

Uzdensky AB (2018) Photothrombotic Stroke as a Model of Ischemic Stroke. Transl Stroke Res 9:437–451.

Virley D, Beech JS, Smart SC, Williams SCR, Hodges H, Hunter AJ (2000) A Temporal MRI Assessment of Neuropathology after Transient Middle Cerebral Artery Occlusion in the Rat: Correlations with Behavior. Journal of Cerebral Blood Flow & Metabolism 20:563– 582 Available at: https://journals.sagepub.com/doi/10.1097/00004647-200003000-00015.

von Bornstädt D, Houben T, Seidel JL, Zheng Y, Dilekoz E, Qin T, Sandow N, Kura S, Eikermann-Haerter K, Endres M, Boas DA, Moskowitz MA, Lo EH, Dreier JP, Woitzik J, Sakadžić S, Ayata C (2015) Supply-demand mismatch transients in susceptible peri-infarct hot zones explain the origins of spreading injury depolarizations. Neuron 85:1117–1131.

Wahl AS, Büchler U, Brändli A, Brattoli B, Musall S, Kasper H, Ineichen B V., Helmchen F, Ommer B, Schwab ME (2017) Optogenetically stimulating intact rat corticospinal tract post-stroke restores motor control through regionalized functional circuit formation. Nat Commun 8.

Wang C, Lin C, Zhao Y, Samantzis M, Sedlak P, Sah P, Balbi M (2023) 40-Hz optogenetic stimulation rescues functional synaptic plasticity after stroke. Cell Rep 42:113475 Available at: https://linkinghub.elsevier.com/retrieve/pii/S2211124723014870.

Wang H, Peca J, Matsuzaki M, Matsuzaki K, Noguchi J, Qiu L, Wang D, Zhang F, Boyden E, Deisseroth K, Kasai H, Hall WC, Feng G, Augustine GJ (2007) High-speed mapping of synaptic connectivity using photostimulation in Channelrhodopsin-2 transgenic mice. Proc Natl Acad Sci U S A 104:8143–8148.

Wang R, Wang H, Liu Y, Chen D, Wang Y, Rocha M, Jadhav AP, Smith A, Ye Q, Gao Y, Zhang W (2022) Optimized mouse model of embolic MCAO: From cerebral blood flow to neurological outcomes. Journal of Cerebral Blood Flow and Metabolism 42:495–509.

Watson BD, Dietrich WD, Busto R, Wachtel MS, Ginsberg MD (1985) Induction of reproducible brain infarction by photochemically initiated thrombosis. Ann Neurol 17:497–504.

Weir P, Maguire R, O’Sullivan SE, England TJ (2021) A meta-analysis of remote ischaemic conditioning in experimental stroke. Journal of Cerebral Blood Flow and Metabolism 41:3–13.

Wester P, Watson BD, Prado R, Dietrich WD (1995) A Photothrombotic ‘Ring’ Model of Rat Stroke-in-Evolution Displaying Putative Penumbral Inversion. Stroke 26:444–450 Available at: https://www.ahajournals.org/doi/abs/10.1161/01.STR.26.3.444.

White AJ, Chintada L, Chan HH, Fisher BM, Mandava N, Hogue O, Machado AG, Baker KB (2025) Sex-specific cell death mechanisms in rodent sub-acute and chronic stroke. Neurosci Lett 863:138307 Available at: https://linkinghub.elsevier.com/retrieve/pii/S0304394025001958.

Yang J, Serrano P, Yin X, Sun X, Lin Y, Chen SX (2022) Functionally distinct NPAS4-expressing somatostatin interneuron ensembles critical for motor skill learning. Neuron 110:3339–3355.e8.

Yin X, Jones N, Yang J, Asraoui N, Mathieu M-E, Cai L, Chen SX (2021) Delayed motor learning in a 16p11.2 deletion mouse model of autism is rescued by locus coeruleus activation. Nat Neurosci 24:646–657.

Yoguim MI, Grandini GS, Bertozo L de C, Caracelli I, Ximenes VF, de Souza AR (2022) Studies on the Interaction of Rose Bengal with the Human Serum Albumin Protein under Spectroscopic and Docking Simulations Aspects in the Characterization of Binding Sites. Chemosensors 10.

Yoon KC, You IC, Kang IS, Im SK, Ahn JK, Park YG, Ahn KY (2007) Photodynamic Therapy with Verteporfin for Corneal Neovascularization. Am J Ophthalmol 144.

Zhang F, Wang LP, Boyden ES, Deisseroth K (2006) Channelrhodopsin-2 and optical control of excitable cells. Nat Methods 3:785–792.

Zhang Q, Wu JF, Shi QL, Li MY, Wang CJ, Wang X, Wang W, Wu Y (2019) The neuronal activation of deep cerebellar nuclei is essential for environmental enrichment-induced post-stroke motor recovery. Aging Dis 10:530–543.

Zhang S, Zhang X, Zhong H, Li X, Wu Y, Ju J, Liu B, Zhang Z, Yan H, Wang Y, Song K, Hou ST (2022) Hypothermia evoked by stimulation of medial preoptic nucleus protects the brain in a mouse model of ischaemia. Nat Commun 13.

Zhang SY, Jeffers MS, Lagace DC, Kirton A, Silasi G (2021) Developmental and interventional plasticity of motor maps after perinatal stroke. Journal of Neuroscience 41:6157–6172.

